# Lipophilic molecular rotor to assess the viscosity of oil core in nano-emulsion droplets

**DOI:** 10.1101/2024.09.25.614997

**Authors:** Mohamed Elhassan, Carla Faivre, Halina Anton, Guillaume Conzatti, Pascal Didier, Thierry Vandamme, Alteyeb S. Elamin, Mayeul Collot, Nicolas Anton

**Author notes:** To whom correspondence should be addressed: Nicolas Anton and Mayeul Collot.

## Abstract

**Hypothesis:** Characterization of nanoscale formulations is a continuous challenge. Size, morphology and surface properties are the most common characterizations. However, physicochemical properties inside the nanoparticles, like viscosity, cannot be directly measured. Herein, we propose an original approach to measuring dynamic viscosity using a lipidic molecular rotor solubilized in the core of nano-formulations. These molecules undergo conformational changes in response to viscosity variations, leading to observable changes in fluorescence intensity and lifetime, able to sense the volume properties of dispersed nano-domains.

**Experiments:** The lipophilic molecular rotor (BOPIDY derivatives) was specifically synthesized and characterized as oil viscosity sensing in large volumes. A second part of the study compares these results with rBDP-Toco in nano-emulsions. The objective is to evaluate the impact of the formulation, droplet size and composition on the viscosity of the droplet’s core.

**Findings:** The lipophilic rotor showed a universal behavior whatever the oil composition, giving a master curve. Applied to nano-formulations, it discloses the viscosity in the nano-emulsion droplets, enabling the detection of slight variations between reference oil samples and the nano-formulated ones. This new tool opens the way to the fine characterization of complex colloids and multi-domain nano and micro systems, potentially applied to hybrid materials and biomaterials.

## 1. Introduction

Molecular rotors represent a particular category of viscosity-sensitive fluorescent compounds that serve as valuable probes for microenvironment characterization, particularly those which are inaccessible through conventional bulk rheological techniques [1]. The term “molecular rotor” refers to small synthetic fluorophores whose fluorescence emission is sensitive to the viscosity of the surrounding environment [2]. The sensing of the microenvironment has become an interesting research area since the local environment is the most relevant factor governing the physical and chemical behavior of surrounding molecules [3]. Fluorescence sensing techniques have played a crucial role in characterizing various properties such as viscosity, polarity, local acidity/basicity, pH, and temperature, thereby contributing to a better understanding of specific location properties [4,5]. Among various biophysical parameters, viscosity sensing plays an important role in many fields [6,7] including biological processes [8], lipid bilayers [9,10], and living cells [11,12].

On the other hand, while measuring the rheological properties of a macroscopic sample is trivial, determining the microviscosity within micro/nano meter-sized objects remains extremely challenging [13–15].

For example, in Ref. [16], the authors described an interesting approach using cobalt ferrite nanoparticles (NPs) to sense the viscosity inside oil-in-water emulsions (coated with oleic acid) or in the continuous phase (coated with PEG), in order to compare with macroscale viscosity. In this case, NPs sizing around 14 nm were used and the micro-rheology obtained from the measurements of their magnetic susceptibility. A second literature example of particle tracking approach was described in Ref. [17], using fluorescent beads of 0.75 µm. In conventional particle-tracking microrheology, the particle’s motion is monitored, and the properties of the surrounding is obtained from the particle’s trajectory. The mean-squared displacements of the beads were followed as a function of the lag time, to obtain the diffusion coefficient, and deduce the medium viscosity. This is an main phenomenological difference between this approach and molecular rotors that undergoes conformational changes.

Particle tracking is an attractive methodology that allows measuring physicochemical properties of dispersed systems. It is noteworthy that the particle-tracking method not only enables the measurement of viscosity but also allows the characterization of viscoelastic properties when external forces are applied to the particles, resulting in a specific response. This phenomenon is referred to as micro-viscoelasticity. When particles are sufficiently large to be responsive to external stimuli, the behaviors and properties of molecular rotors remain unaffected by such influences. Consequently, molecular rotors can be employed in a similar capacity to particle-tracking approach in a force-free environment, with the distinction that they operate on a different scale due to their smaller size. Behind the fact that this method needs specific experimental setup, the main limitation indeed lies in the scale range of the domains in which they allow to measure the viscosity. This scale range is, in the best condition, a few orders of magnitudes higher than the particles used, and finally will limit micro domains to micrometer size. In contrast, molecular probes (like molecular rotors) will not have these limitations and will allow measurements in domains with much lower scale, up to nano-domains.

Other methods were reported in literature, we can cite the use of diffusing wave spectroscopy, Ref. [18] which can be considered as an implemented dynamic light scattering. This approach allows assessing micro-rheology data from dispersed samples such as ageing, stability, and structural properties, in complementary to macro-rheology. Although this method is simple and interesting, it cannot be applied to study the internal properties of dispersed systems, all the more so since in the nanoscale. We can finally cite another experimental setup based on colloidal-probe AFM, in Ref. [19]. In that configuration, the micro-scale sample is set up between an oscillating tip and a support, to investigate the interfacial mechanics and dynamic rheological properties. The data collected are, to some extent, comparable to those obtained with macro-rheology. Here also, the main difference with the molecular probe is the scale of measurement (not transposable to nanoscale), and the additional constraint to confine the sample onto an AFM tip.

These studies not only emphasize the need and interests in the development of new approaches to measure the fine physicochemical properties of dispersed micro and nano-systems, but also the real innovation and interest to develop new experimental approaches dedicated to nano-scaled colloids in general, thus the originality of our methodology with molecular rotors.

Molecular rotors offer a convenient means of measuring microviscosity by undergoing changes of intramolecular rotation and non-radiative relaxation, thus leading to observable changes in fluorescence intensity and lifetime. The rate of these conformational motions is directly influenced by viscosity. Therefore, variations in local viscosity induce significant changes in both fluorescence lifetime and fluorescence intensity [20]. Consequently, higher local viscosity restricts conformational changes of the molecular rotor, resulting in increased fluorescence lifetime and intensity. It is noteworthy that the study of fluorescence lifetime offers numerous advantages for assessing viscosity over fluorescence intensity; it is independent of the dye concentration. Also, it appears to be one of the most reliable and accurate methods for evaluating viscosity by using molecular rotors, eliminating potential experimental errors [21–23]. In addition, fluorescence lifetime detection can be integrated with imaging techniques like fluorescence lifetime imaging (FLIM) [24].

In this study, we used BODIPY-based molecular rotor to measure the viscosity in the core of lipid nano-emulsions (NEs). NEs are lipid-oil droplets stabilized with surfactants, typically ranging in size from 20 to 300 nm [25,26]. Due to their stability and biocompatibility, NEs are emerging as promising carriers in various fields, including drug delivery, diagnostics, cosmetics, pesticides, and the food industry [27–30]. Compared to other nano-carriers, NEs have garnered attention as bio-mimicking “green” nanocarriers with significant potential for preparing contrast imaging agents and nanomedicines [31]. NEs have the distinct advantage of serving as liquid reservoirs for lipophilic active pharmaceutical ingredients (APIs) and/or lipophilic fluorescent probes molecules, dispersed in aqueous medium [32,33]. This characteristic allows NEs to encapsulate a large number of fluorescent dyes with reduced self-quenching, resulting in high quantum yield and ultrabright fluorescent nanomaterials. The brightness of these nano-droplets opens up new possibilities, such as single droplets tracking in cells or *in vivo* in small laboratory animals [34–36].

To increase the encapsulation efficiency of fluorophores in NEs they should be highly soluble in oil to achieve high dye loading; sufficiently bulky to prevent aggregation-caused quenching (ACQ) [37], which would reduce the quantum yield of dyes encapsulated. Additionally, these dyes should be hydrophobic enough to remain within the NEs and photostable enough to enable tracking over time [34]. However, various parameters, such as hydrophobicity, polarity, composition, solubility, temperature, and viscosity, can significantly influence not only the physicochemical properties of the NEs but also the photophysical properties of the encapsulated fluorophores, leading to significant changes in the fluorescence properties of the NEs [36,38]. To date, the literature on nano-emulsions –and more generally on emulsions– considers that the oil core of formulated droplets conserves similar properties as the native oil phase. This affirmation could, indeed, be considered as a corollary of the Bancroft’s rule [39,40]. In general, the formulation of emulsions is considered producing oil droplets stabilized with a layer of stabilizers in the interfacial region. These amphiphiles molecules are soluble in the continuous phase, which results in the sense of the interfacial curvature. This is finally why we generally neglect their interactions with the droplet composition.

Fluorogenic molecular rotors (FMRs) have demonstrated their efficiency in probing viscous environments through a linear correlation between fluorescence intensity, or lifetime, and viscosity. Green-emitting, and non-charged dyes known as 4,4-difluoro-4-bora-3a,4a-diaza-s-indacenes (BODIPYs) have become widely used in bioimaging. Indeed, these rotors serve to label proteins and DNA due to their high rotational ability and brightness [41]. This family of fluorescent probes is also used to characterize the modification of their solubilizing medium, as it is potentially linked to their fluorescence properties. However, understanding the relationship between the physicochemical properties of the environment in which the dyes are solubilized, the formulations, and the resulting fluorescence efficiency remains limited. To this end, we proposed to evaluate the relationship between the viscosity and fluorescence of rBDP-Toco, a derivative of BOPIDY, which is conjugated with α-tocopherol to enhance the rotor hydrophobicity [42], and allowing an efficient solubilization in oils. This BODIPY rotor moiety used in this study is a golden standard and was abundantly characterized in the literature, notably by the group of Kuimova *et al.* [1], and was also applied to bioimaging in previous reports [43,44]. The advantage of this BODIPY rotor is its high sensitivity toward variations of viscosity notably using fluorescence lifetime and its low sensitivity toward other environmental changes like temperature [45] or polarity [46].

In the present study, we render the well-known molecular rotor lipophilic by coupling it with a lipid moiety (tocopherol), as illustrated below (scheme 1) that presents the nano-emulsion formulation. We have previously shown that, when coupled to BODIPY, tocopherol drastically enhanced the BODIPY solubility [42]. This coupling does not affect the viscosity sensitivity, nor it modifies the fluorophore structure. In addition, it is important to note that this lipophilic BODIPY molecular rotor is recognized to have a very weak sensitivity to solvent polarity, as described in previous studies [46].

**Scheme 1:**
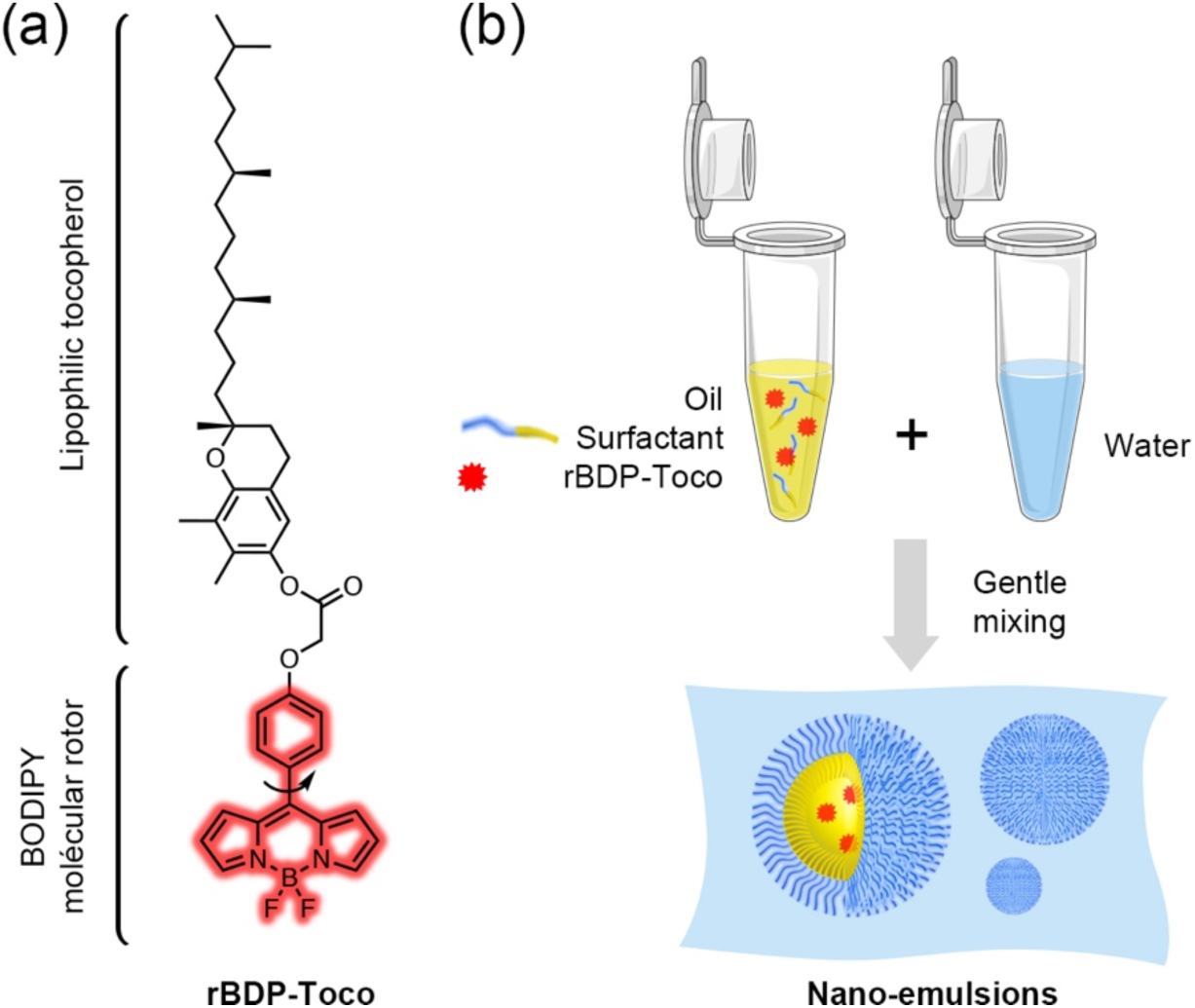
(a) Structure of rBDP-Toco. The viscosity fluorescent reporter is a BODIPY molecular rotor, the lipophilicity was enhanced by coupling with the lipid tocopherol. (b) Principle of the nano-emulsion formulation by spontaneous emulsification.

The first part of this study focuses on characterizing the lipophilic molecular rotor, in order to establish the correlation between the actual viscosity of the oily phase of pure oil mixtures and optical properties. The second part is dedicated to the characterization of the oil properties when in nano-formulation by comparing the actual viscosity with the viscosity of the oil core. We emphasize the potential difference arises between fluorescence characterization –generally adopted in the reports on nano-emulsions– and fluorescence lifetime that only focus on the molecular behavior independently to concentration issues. However, the simultaneous influence of temperature on viscosity and the photophysical behavior of fluorophores is often overlooked [47]. Nevertheless, temperature variations can introduce significant offset in viscosity readouts, and vice versa. Therefore, the final part of this study investigates the effect of temperature on the photophysical behavior of rBDP-Toco.

To conclude, this study proposes a new approach to sense the properties of nano-scaled formulation, and potentially applicable to a broad range of materials, nanomaterials and multiscale materials in general. For this reason, it presents a high potential if the characterization of the colloids, and in the understanding of their behavior and composition.

## 2. Materials and methods

### 2.1. Materials

The oil phase, vitamin E acetate (VEA), was provided by Tokyo Chemical Industry (Tokyo, Japan), castor oil from Sigma-Aldrich (France), and Labrafac WL 1349 (medium chain triglycerides, MCT) was obtained from Gattefossé (Saint-Priest, France). The nonionic surfactant used, Kolliphor® ELP was purchased from BASF (Ludwigshafen, Germany). A stock solution of 4.3 mM of the rBDP-Toco dye in dioxane was prepared beforehand (structure of rBDP-Toco is reported in Scheme 1 (a)). Milli-Q water was obtained from a Millipore filtration system and used in all experiments. All chemicals were of analytical grade. rBDP-Toco was synthesized, the protocols and characterizations can be found in the supplementary information.

### 2.2. Methods

#### 2.2.1. Preparation of the oils mixture

In order to prepare oils possessing increasing viscosities, highly viscous oils (VEA and castor oil) were mixed at various ratios with lower viscosity MCT, thus we prepared: *(i)* MCT/castor oil and *(ii)* MCT/VEA. The oil mixture was homogenized at 80°C in a thermomixer (Eppendorf) for 5min minutes, and then vortexed for 5 minutes. The different oil ratios were gradually modified, as described in the *results* part below, with a ratio of the more viscous oil varying from 0 wt.% wt. to 100 wt.%.

#### 2.2.2. Rheological characterization

Rheology was performed on a HAAKE MARS 40 Rheometer (Thermo Scientific) using geometries of 35 mm diameter parallel plates with a gap of 0.1 mm. The volume of the oil sample (0.1 mL) was added to the plate geometry, and the temperature was set at 20°C. The rotational rheometer was in *controlled rate* mode for which a shear rate (γ̇) was applied and shear stress (τ) measured. The applied range of γ̇ was from 0.001 to 1000 s^-1^. All the measurements were performed in triplicate.

#### 2.2.3. Absorption and fluorescence spectra

Absorbance and fluorescence spectra of oil mixtures were performed with a Thermo Scientific™ Varioskan™ LUX multimode microplate reader. The absorption spectra were scanned from 300 to 700 nm, while the emission spectra were recorded from 478 to 700 nm with an excitation wavelength of 460 nm. The sample prepared for analysis was a mixture of 4.6 µL of the rBDP-Toco dye (of stock solution at 4.3 mM in dioxane) and 1995.4 µL of oil. Then, 150 µL of the prepared sample was analyzed. Transparent microplates were used for absorbance spectra analysis and black microplates for fluorescence analysis. Fluorescence intensities of nano-emulsions were measured according to the same methodology, and, in order to compare their values, each other, FI are normalized with oil amount as reference (as oil amounts can vary between the different formulations and pure oil mixtures). All the measurements were performed in triplicate.

#### 2.2.4. Fluorescence Lifetime Measurements

Time-resolved fluorescence measurements were performed with the time-correlated single-photon counting technique with excitation at 500 nm (supercontinuum laser NKT Photonics SuperK Extreme with 10 MHz repetition rate). The fluorescence decays were collected at 515 nm using a polarizer set at magic angle and a 16 mm band-pass monochromator (Jobin Yvon). The single-photon events were detected with a micro-channel plate photomultiplier R3809U Hamamatsu, coupled to a pulse pre-amplifier HFAC (Becker-Hickl GmbH) and recorded on a time-correlated single photon counting board SPC-130 (Becker-Hickl GmbH). Time-resolved exponential decays were fitted by using a function corresponding to a exponential decays convolved with a normalized Gaussian curve of standard deviation σ standing for the temporal IRF and a Heavyside function. The fitting function was built in Igor Pro (Wavemetrics). All emission decays were fitted using a weighting that corresponds to the standard deviation of the photon number squared root. The lifetimes were measured for the pure oil mixtures (castor oil/MCT and VEA/MCT mixtures) with different viscosities to establish the calibration of the molecular rotors with actual volume viscosity obtained with the oscillatory rheometer.

#### 2.2.5. Formulation and Characterization of Nano-emulsions

Nano-emulsions (NEs) were formulated by the spontaneous emulsification method. As illustrated in Scheme 1 (b), the first phase of {oil + nonionic surfactants} is heated and homogenized, and then suddenly mixed with aqueous phase (MilliQ water) at 80°C. As a result, the oil phase is broken up and generates oil-in-water nano-emulsion droplets [48]. Loading such nano-emulsion with a dye only consists in solubilizing the probe in the oil phase and follow the same formulation process, without impact on the droplet’s properties. The proportions with the nonionic surfactant Kolliphor ELP® were defined as surfactant-to-oil weight ratio, defined as SOR = 100 × *w*_surfactant_/ (*w*_surfactant_ + *w*_oil_) [49], where *w* is the weight of the different compounds, varied from 50% to 70%.

#### 2.2.6. Characterizing nanodroplet size distribution

The droplet size distribution, mean size, and polydispersity indexes (PDIs) were determined by dynamic light scattering (DLS) [50] using a Malvern^®^ Nano ZS instrument (Malvern, Orsay, France). The nano-emulsion samples were diluted 10 times in distilled water. Size distribution and polydispersity index (PDI) were recorded at a temperature of 25 °C. All the measurements were performed in triplicate.

#### 2.2.7. Statistical analysis

All the quantitative data are expressed as mean +/-standard deviation obtained from three independent experiments.

## 3. Results and discussion

### 3.1. Rheological characterization

Dynamic viscosities (η) are represented against shear rate γ̇, and report on *supplementary information* (Fig. S1) for the different oil mixtures. η shows a gradual decrease and plateau stabilization, which is actually a phenomenon known for lower values of shear rates with lower intermolecular interactions [51], and stabilize at higher values of shear rates to a Newtonian behavior. In this study, we collect the values of stabilized viscosities in the plateau region (η_∞_), which are reported in Fig. 1, for the two oil mixtures. The number of measurements was adapted to the oil mixtures, with a higher number for VEA / MCT mixture since the curve underwent a higher variation in the 80% – 100% range.

**Figure 1:**
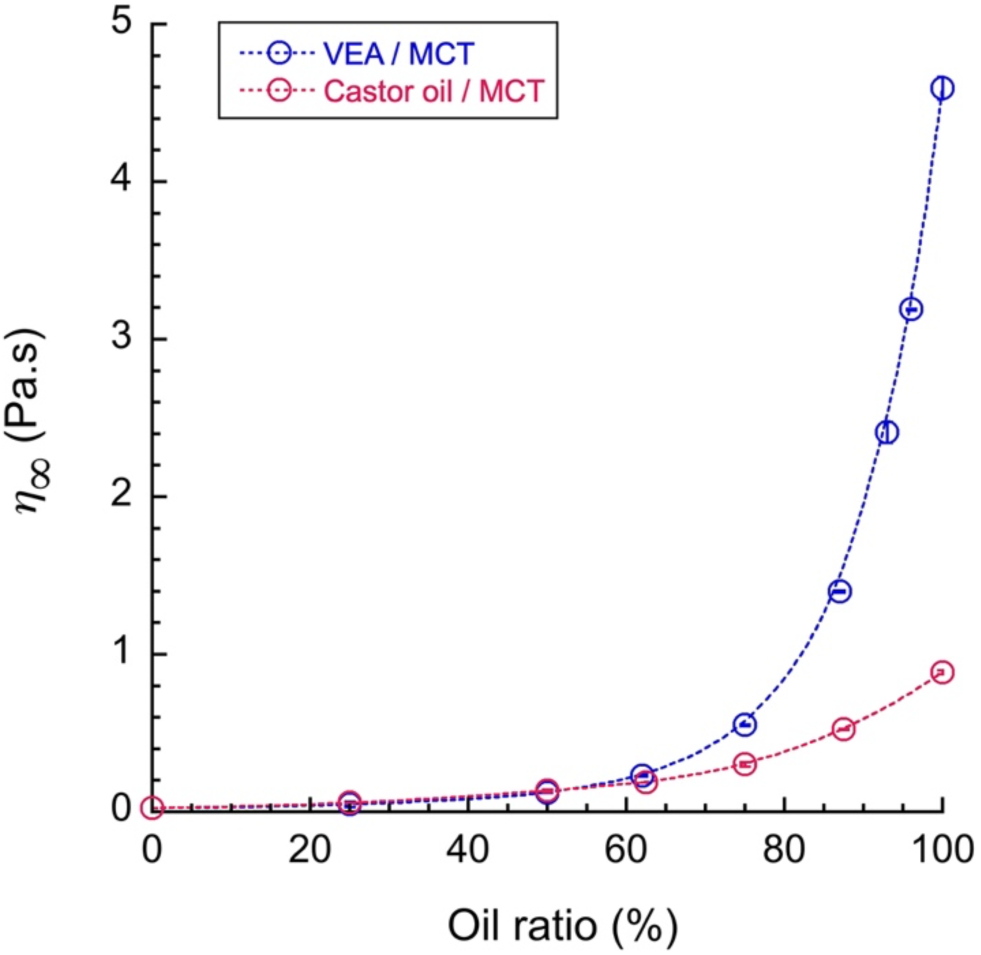
Dynamic viscosities of oil mixtures. VEA or castor oil concentration (%) mixed with MCT.

For pure MCT, η_∞_ tends to (24.0 ± 0.1) mPa.s, and the two curves are superimposed up to about 60%, then show a drastic separation that allows a significant difference in the properties of the two oil mixtures.

### 3.2. Fluorescence characterization of rBDP-Toco

The molecular rotor rBDP-Toco is dissolved in the different oil mixtures, on which a characterization of the fluorescence properties is performed. It shows for each oil mixture a maximum peak at 512 nm. On the other hand, once excited at 460 nm, fluorescence intensities (FI) are measured at 512 nm, and their values are reported in Fig. 2 (a) in function of the oil composition (oil ratio of VAE and castor oil in MCT), and in Fig. 2 (b) as a function of their viscosities obtained from Fig. 1.

**Figure 2:**
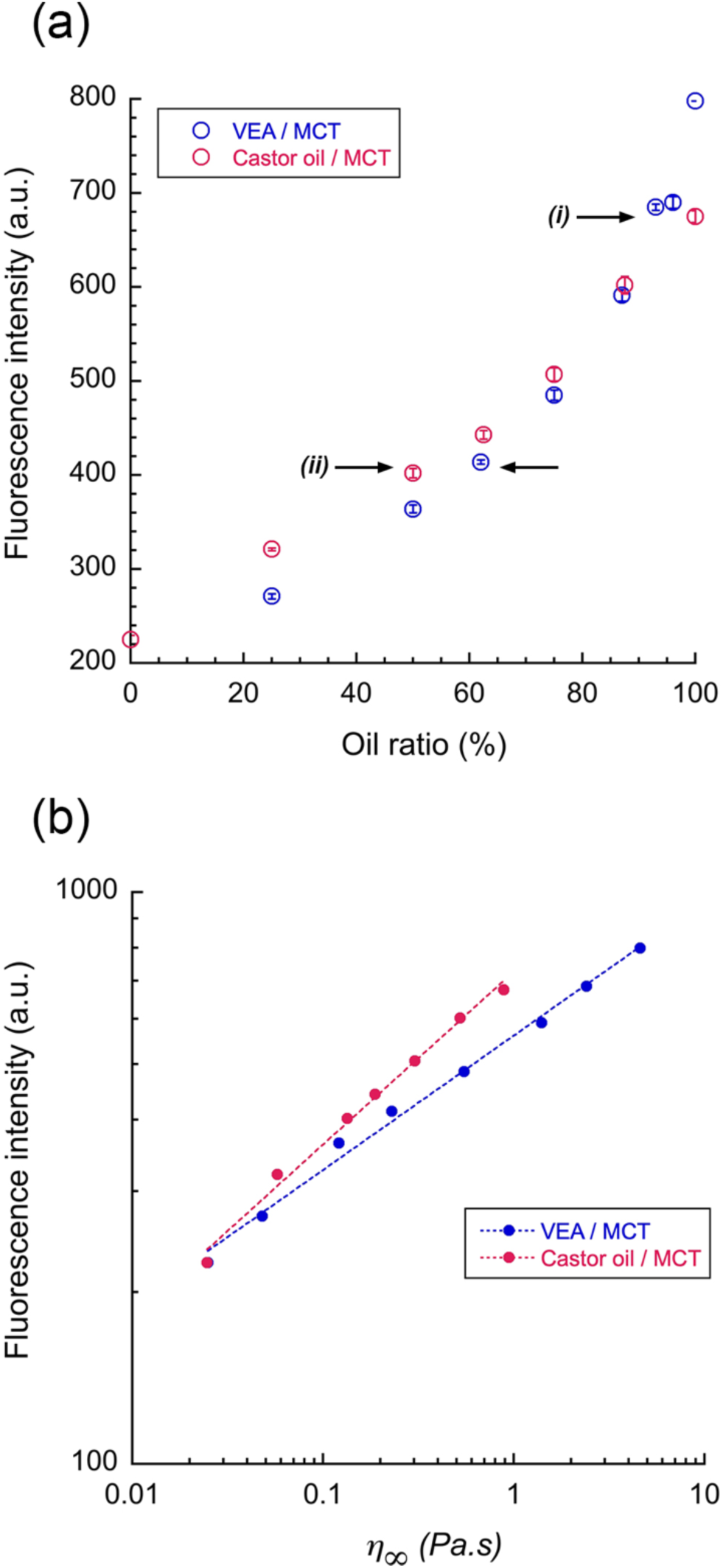
Fluorescence intensity (λ_*ex.*_ = 460 nm, λ_*em.*_ = 512 nm) of rBDP-Toco dissolved in different oil mixtures, represented (a) against VEA or castor oil concentration (%) mixed with MCT, and (b) against the corresponding viscosities of these oil mixtures (obtained from Fig. 1).

Fig. 2 (a) shows first the correlation between FI of rBDP-Toco, with the viscosity of its solubilizing medium. As the viscosity of the medium increases, the fluorescence intensity of the fluorogenic molecular motor increases. This can be explained by the restriction of intramolecular rotation commonly observed in fluorophores with the structure of molecular rotors, a phenomenon known as ‘motion-induced change in emission’.

On the other hand, Fig. 2 (a) compared oil mixtures with different viscosities and reveals very similar FI values –whereas clear difference has been expected. When FI values are transposed in function of η_∞_, in Fig. 2 (b), two distinct curves are disclosed, described by power laws. It follows therefore that rBDP-Toco has a clear response to a viscosity rise, but the efficiency of this response depends on the nature of oil. For instance, if we consider the FI⁄η_∞_ ratio for representative points with similar FI, *e.g.* for FI ≈ 680 a. u. and FI ≈ 400 a. u. (shown as arrows *(i)* and *(ii)* in Fig. 2 (a)), FI⁄η_∞_ appears significantly different. For example *(i)*, FI⁄η_∞_ appears equal to 284 a.u.Pa^-1^.s^-1^ for VEA mixture and 760 a.u.Pa^-1^.s^-1^ for castor oil mixture, while for the case *(ii)*, FI⁄η_∞_ appears equal to 1800 a.u.Pa^-1^.s^-1^ for VEA mixture and 2990 a.u.Pa^-1^.s^-1^ for castor oil mixture.

These first results emphasize significantly different fluorescent behavior of rBDP-Toco in function of its solubilization medium, and thus the limitation of this approach to be considered as a universal viscosity probe. This result is likely due to the slightly different efficiency of these oils to solubilize such a molecular rotor, which can result in slight differences in the actual dye concentrations in oil.

Consequently, fluorescence lifetimes (FLT) were measured for all these samples. The main difference from the simple fluorescence intensity is that lifetime measurement is not sensitive to concentration or related solubility issues. Results are reported in Fig. 3, and as opposed to those of FI, the FLT appears independent of the oil’s composition and the two oil mixtures show results that can be superimposed.

**Figure 3:**
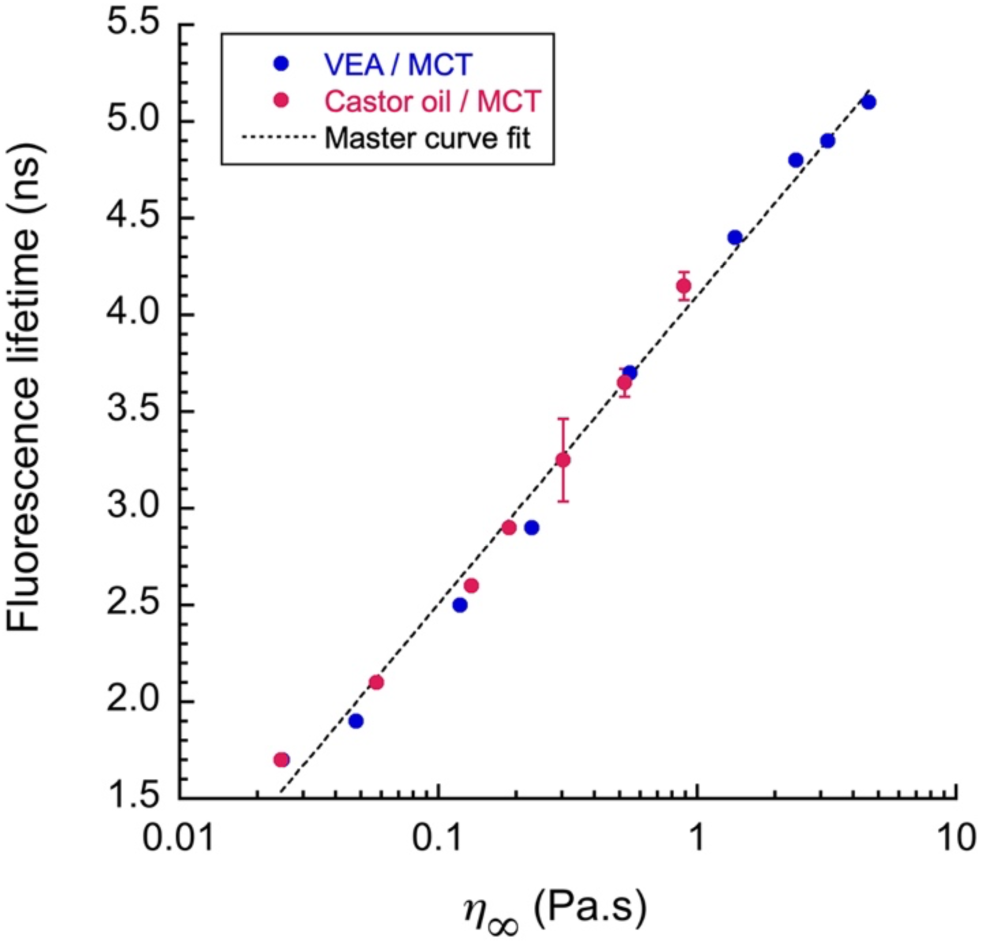
Fluorescence lifetime (λ_*ex*._ = 460 nm, λ_*em*._ = 512 nm) of rBDP-Toco dissolved in different oil mixtures, represented against viscosities of oil mixtures of VEA or castor oil concentration (%) mixed with MCT.

This result is critical and shows the potential of such molecular rotors as viscosity probe independently to the nature of the oil. When fluorescence characterization is classically associated with the measurements of FI, thus we can consider FLT here as the reference. However, comparing FLT and FI remains interesting to refine the characterization of the probe in its solubilizing medium: in the present case, rBDP-Toco appears more soluble in MCT/castor oil mixtures compared to MCT/VEA ones. The logarithmic fit gives the following calibration curve of rBDP-Toco: FLT = 4.1 + 1.6 × log (η_∞_) with a coefficient of determination equal to R²= 0.996.

A complementary set of experiments was undertaken, to investigate the effect of temperature on the behavior of rBDP-Toco dye, and reported in supplementary materials as Fig. S2. Viscosity and FLT were measured for VEA / mixture over a temperature range from 20°C to 55°C, (Fig. S2 (a) and (b)) for representative ratio (VEA ratio from 75% to 100%) and represented together (Fig. S2 (c)). This analysis showed that the FLT recorded at the same temperature did not overlap, and as well, were not superimposing with the master curve at 20°C. This finding suggests that the behavior of the molecular rotor is strongly dependent on the temperature and does not allow comparing different compositions at different temperatures. Fig. S2 (c) confirms that changing the study temperature would alter the master curve, the complex interplay between these factors, and emphasizes the importance of considering both viscosity and temperature effects in studying such systems. In summary, it appears that, at a given temperature, molecular rotors are accurate viscosity sensors, for different types of lipophilic phases, and can even reveal slight variations in their composition. However, changing the temperature reduces viscosity and FLT, but far from the master curve established. Thus, we can consider that different master curves can be established for each different temperature.

### 3.3. Measuring the viscosity in nano-emulsions

In order to understand the properties of nano-emulsions and the impact of the nanoscale formulation on the oil core of the droplets, we formulated the aforementioned different oil mixtures as nano-emulsions. The properties of nano-emulsions formulated by spontaneous method –*i.e.* size and dispersity– depend on the nature of the oil and the amount of surfactant used. The higher the surfactant amount, the lower the nano-droplet size [52]. Nano-emulsification occurs in a specific condition when oil, surfactant and water are mixed together, and widely described [25]. However, the efficiency of the spontaneous emulsification process depends on the relative affinities of the surfactants with oil and water, and thus with slightly differ when using different oils. In this study, we have selected 3 different surfactant amounts and followed the size and polydispersity for the two oil mixtures. The results are reported in Fig. 4 and emphasize the respective effects of the different oils in the droplet’s size. Representative size distributions are reported the supplementary materials section, Fig. S3. The more efficient process (*i.e.,* the process that gives the smaller droplets) arises when using VEA, giving sizes below 50 nm, then, MCT gives droplets sizing around 100 nm and castor oil produces sizes slightly higher but below 130 nm (Fig. 4 (a) and (b)). The PDI values (Fig. 4 (c) and (d)) are below 0.2, show a good monodispersity of the droplet population and validate the size results.

**Figure 4:**
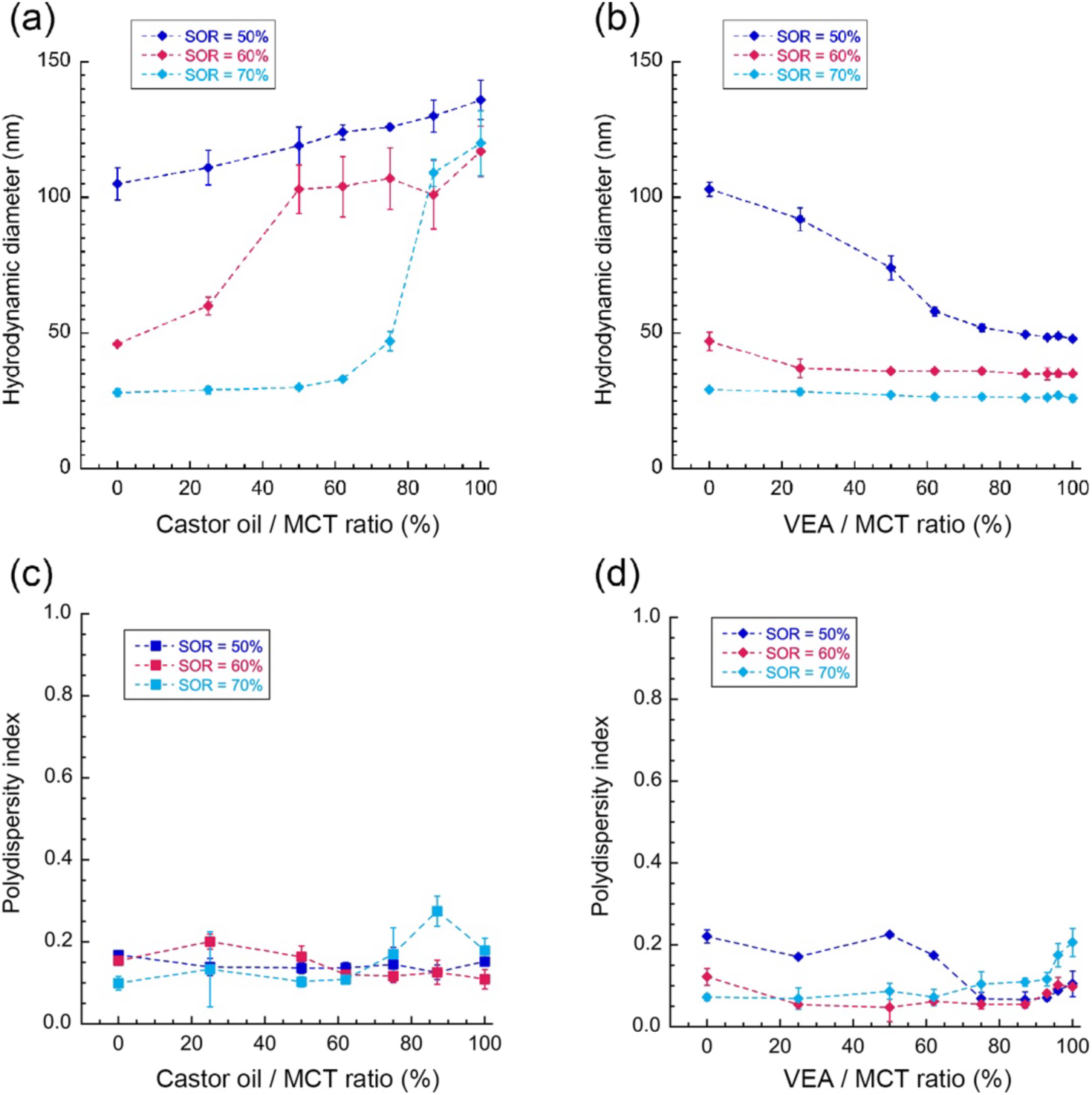
Nano-emulsion characterization. Hydrodynamic diameter (a) and (b), and corresponding PDI (c) and (d), for the different oil mixtures, for three surfactant amounts.

These variations are due to the difference between surfactant affinities for these various oils, which could modify the mobility of surfactants and water in the {oil + surfactant} phase, which results in the spinodal decomposition of the oil phase [53]. This behavior was also reported with change in the oil composition or oil chemical structures [42,54–56]. Interestingly, despite variations in the viscosities of these oils, no clear correlation between viscosity and the results depicted in Fig. 4 is apparent. Even, the more viscous oil VEA appears to allow the most efficient nano-emulsification process by giving the smallest droplets and, while the MCT showing the lowest viscosity gives lower sizes than castor oils. Previous research has provided conflicting findings regarding the influence of oil phase properties on the formation of nano-emulsions (NEs) via spontaneous emulsification. Our observation of a lack of strong correlation between particle size and the physicochemical properties, while it could be correlated only to the chemical structure of oils, which the lower molecular weight (VEA) gives the smaller sizes. Therefore, the physicochemical mechanisms governing the size of droplets produced by spontaneous emulsification remain unclear, warranting further basic research in this area [57]. In addition, the size gradually changes along the composition of the oil mixtures, as seen in Fig. 4 (a) and (b). We can note the regular monodispersity along the different formulations, with PDI generally below 0.2.

Herein we compare the viscosities of pure oils in cuvettes, with the one of the cores of nano-emulsion droplets. Thus, nano-emulsions encapsulating rBDP-Toco were formulated, with different viscosities –according to the two oil mixtures at different ratios. In this part, our objective was to measure the viscosity of oil when incorporated into nano-emulsions and to compare it to one of the pure oils or oil mixtures used for their formulation —disclosing whether the nano-formulation modifies its value. The fluorescence lifetime was chosen to prevail upon solubility and concentration issues (as compared to fluorescence intensity values). To this end, FLT values were compared to the master curve established above in Fig. 3, which is reported in Fig. 5, where two different surfactant amounts –and size ranges– are compared. Fig. 5 (a) shows larger nano-emulsions (made with the lower surfactant amount, SOR = 50%) and Fig. 5 (b) shows smaller nano-emulsions (made with the higher surfactant amount, SOR = 70%).

**Figure 5:**
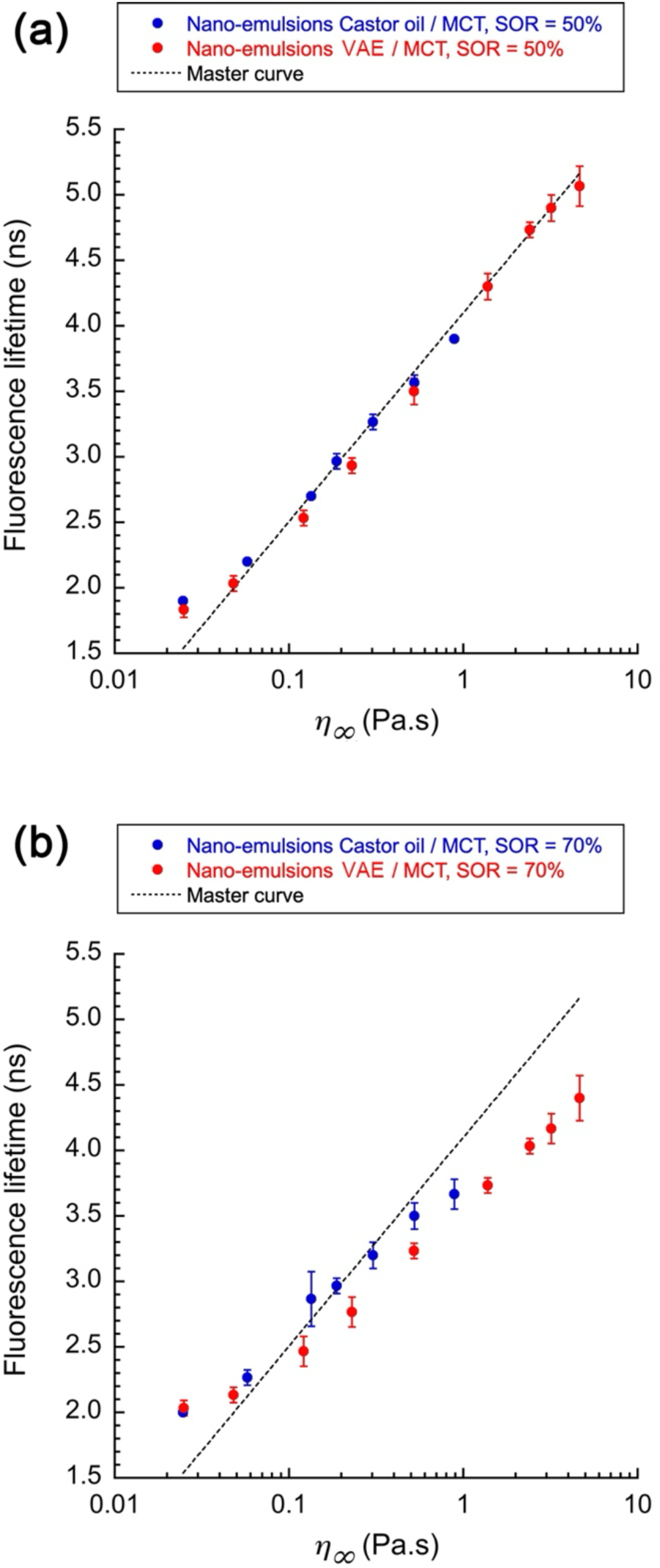
FLT of nano-emulsions made with different oil mixtures (detailed in the legend), as a function of the viscosities (η_∞_) of the corresponding oil mixtures composing their core (from Fig. 1). (a) SOR = 50% and (b) SOR = 70%; n = 3.

Two behaviors are disclosed: *(i)* the exact superimposition of the experimental point and the master curve for SOR 50% nano-emulsions, and *(ii)* the slight shifting between the two curves for SOR 70% nano-emulsions. In case *(i)* we can deduce an exact correspondence between the viscosities of pure oil and oil in the core of the nano-droplets, whereas, in the case *(ii)*, the slight shift is due to a difference between their viscosities.

In order to investigate in detail this difference, Fig. 6 reported the comparison of the viscosities of nano-emulsion droplets noted η_app_(and calculated from the master curve with FLT), with the ones of pure oils (η_∞_). This comparison appears linear, with the slope *s*, giving an idea of the real viscosity of the oil within the nano-droplets, which appears to be decreased compared to pure oil mixtures.

**Figure 6:**
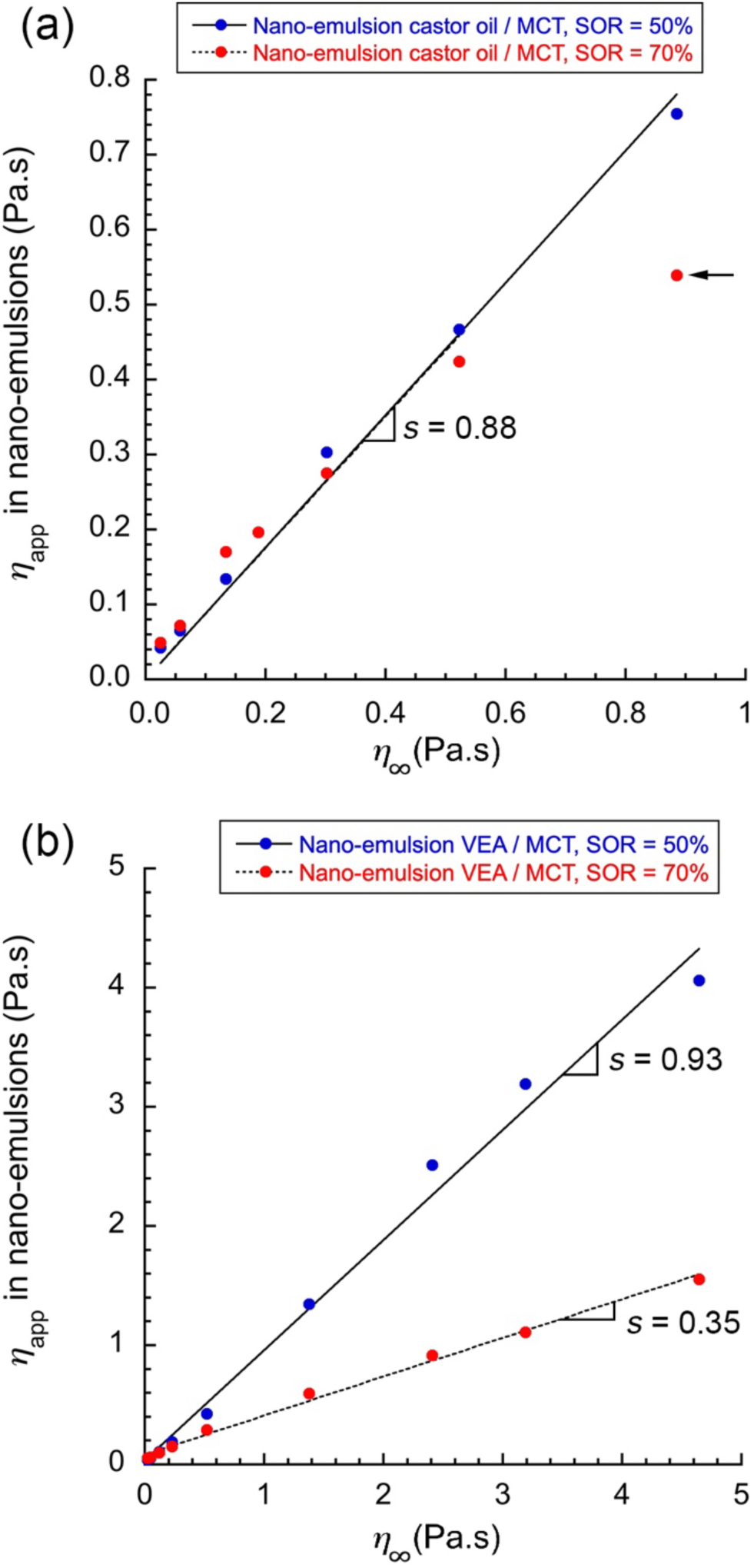
Comparison of viscosities between oil mixture in the core of nano-emulsion droplets (η_app_) and pure oil mixtures (η_∞_), for two different SOR (SOR = 50% and 70%), and for two different compositions of oil mixtures: castor oil / MCT (a), and VEA / MCT (b). Linear extrapolation and related slopes (*s*) are indicated in the figures (excluding nano-emulsions formulated with pure castor arrow).

In the case of castor oil / MCT mixtures (Fig. 6 (a)), nano-emulsion droplet’s core shows a viscosity very similar to the two formulations (SOR 50% and 70%), with a slope *s* = 0.88 exactly equal to SOR = 50% and SOR = 70%. It follows therefrom that nano-formulation would have a viscosity of around 90% of the pure formulation, and this difference would be the direct consequence of its intrinsic composition, including a part of the surfactants remaining mixed with oil in the droplets’ core. Indicated by the arrow in the figure, nano-emulsions formulated with pure castor oil were remarkably deviated from the general trend with a viscosity significantly lower than pure oil.

As regards the VEA / MCT mixture, a clear difference between the two types of nano-emulsions was revealed, with a slope *s* = 0.93 for SOR = 50% that reaches a value of *s* = 0.35 for SOR = 70%. This interesting result signifies that increasing the surfactant amount results in a significant drop in the droplets’ core viscosity or around 35% of the initial value of pure oil mixture –and whatever the ratio of VEA / MCT. This decrease occurs in specific conditions, particularly when using VEA and large surfactant amounts. Therefore, it can be explained by the presence of nonionic surfactants in the oil core of the droplets, which impacts its composition and viscosity. Here this result refers to a particular case of SOR = 70%, with nano-emulsions made with VEA / MCT, where the apparent viscosity η_app_ in nano-emulsion is much lower than the same pure oil mixture, and our hypothesis here is a modification of the oil composition by the surfactants. It is interesting to note that these results do not appear correlated to the change in the droplet’ size reported in Fig. 4. For instance, at SOR = 50%, the VEA / MCT mixtures the underwent a drastic droplet drop along with a FLT close to the behavior in cuvette. When SOR = 50%, the surfactant amount is not sufficient to impact the core properties, which appears similar to the corresponding oil phase –around 93% of the native viscosity– and when SOR = 70% it appears significant to impact the oil composition. On the other hand, this phenomenon is not visible with the castor oil / MCT mixture, even at the highest surfactant amount. In that case, we can consider the surfactant is not solubilized within the droplets’ core. This result refers to the case of SOR = 70%, with nano-emulsions made with castor oil / MCT, were the apparent viscosity η_app_in nano-emulsion is comparable to the viscosity of the corresponding pure oil mixture. In that case, we can consider the oil composition (in NEs and in volume) as similar. A possible explanation is the potential change of the oil’s droplet composition as the result of integration of surfactants in oil core of nano-emulsion droplets. As this difference occurs when comparing castor oil and VEA, we can conclude it depends on the nature of the oil, and thus on the oil / surfactant affinity.

In order to investigate the role of nonionic surfactants in the change in the viscosity, and to understand the limits of our hypothesis, we selected a representative point (pure VEA, as opposed as VEA / MCT mixtures), and we measured the FLT of rBDP-Toco in mixtures of VEA with different amounts of nonionic surfactants. The results are reported in Fig. 7, presenting the viscosity η_∞_ as a function of the composition (Fig. 7 (a)) and FLT plotted against the viscosity η_∞_ (Fig. 7 (b)).

**Figure 7:**
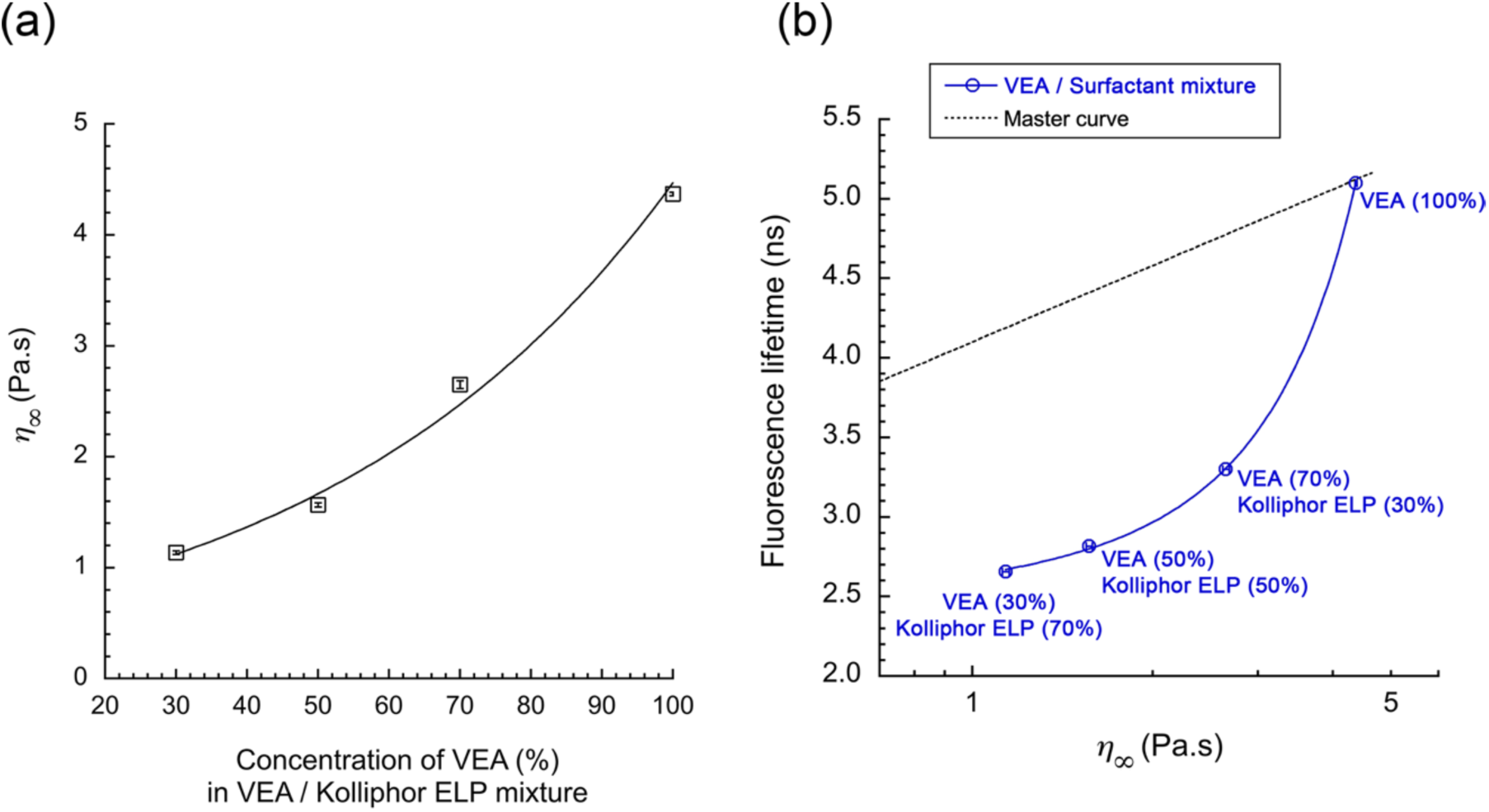
Characterization of the mixture VEA / Kolliphor ELP (nonionic surfactant). (a) Viscosity for different compositions, and (b) FLT of rBDP-Toco as a function of the viscosity, in these VEA / Kolliphor ELP mixtures (where the composition is indicated as labels in the graph). The comparison with the master curve obtained with oil mixtures is also reported in the figure; n = 3.

The first important point (in Fig. 7 (a)) is a significant decrease of η_∞_. That would corroborate the trend observed in Fig. 6 with nano-emulsions (VEA, SOR = 70%), and our hypothesis to attribute this loss of FLT to the viscosity decrease induced by the modification of the droplet composition. However, in Fig. 7 (b), the correlation between FLT and viscosity appears to not follow the master curves established with the oil phases, but another behavior resulting from the presence of nonionic surfactants. FLT decreases much faster compared to the master curve, In contrast with the former experiment with simple oil mixtures, the surfactant / oil mixture is a fundamentally different system, and this experiment was, indeed, conducted to verify our assumption regarding the change of viscosity induced by a potential integration of surfactants in the oil core of emulsion.. Nevertheless, in the case of nano-emulsions made with VEA as oil, and for SOR = 70% (see Fig. 5 (b) above), FLT value is (4.4 ± 0.2) ns. In that case, considering our hypothesis –that the loss of FLT can be due to integration of surfactants in oil– the fitting exponential function (Fig. 7 (b); FLT = 2.19 + 0.25 · *e*^(0.56×η∞)^) allow evaluating the viscosity in nano-emulsions, as η_∞_;^*NE*^ = 3.89 Pa.s. Finally, from the extrapolation of the results given in Fig. 7 (a) (η_∞_ = −0.79 + 1.21 · *e*^(0.014×[VEA])^), we obtain the composition 91.8% of VEA and 8.2% of Kolliphor ELP, in the nano-emulsion core. Considering the SOR of 70% in the composition of the nano-emulsion, one can note that such a proportion seems nevertheless low, with a significant impact on the viscosity. It is possible that the presence of surfactants brings water molecules. However, as discussed above, these lipophilic molecular rotors are not sensitive to the polarity, and the solubility will not affect the FLT. Therefore, we should not see the impact of water on the measurement of viscosity. On the other hand, such modification of the composition can impact on the value of viscosity (as seen in Fig. 7).

To summarize, molecular rotors allow measuring the viscosities in the nano-emulsion droplets cores, and can disclose changes due to a modification of their composition. For very high surfactant concentrations, amphiphilic molecules can be incorporated in the oil core and impact on the photophysical properties of the BODIPY rotor. Therefore, the FLT / viscosity calibration should be carefully adapted to this specific system. It also shows that the nonionic surfactant used in the formulation has a better and specific affinity for VEA acetate compared to MCT and castor oil. Interestingly, this result highlights the strong correlation between the oil/surfactant affinity and the efficiency of the nano-emulsification process; as the droplet’s size in function of the composition (Fig. 4) is much more efficient with using VEA.

In this first series of experiments, the master curve showed a direct correlation between the FLT and oil viscosities –regardless of the composition of oil. On the other hand, fluorescence intensities were comparable between the two different oil mixtures –castor oil / MCT and VEA / MCT. In general, this difference is attributed to the difference in solubilities of the rotor rBDP-Toco in the oil phase, which avoids using FI as a criterion to assess the values of viscosities. Nevertheless, as the composition of oil is also changed with the nano-emulsion formulations, it remains interesting to study the values of FI of nano-emulsion droplets, compared to pure oils. These results are reported in Fig. 8, comparing the two oil mixtures.

**Figure 8:**
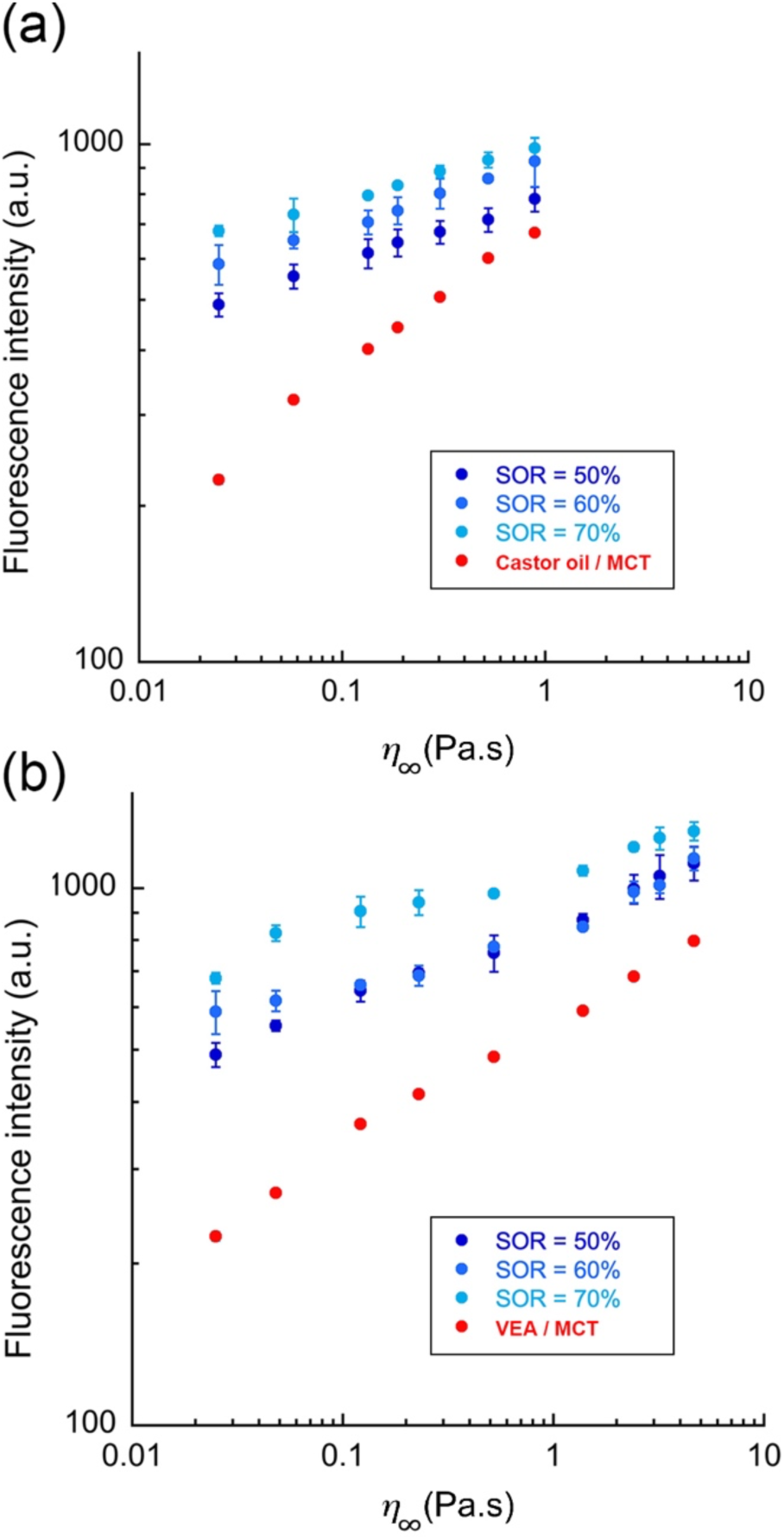
FI of oil mixtures and their respective nano-emulsions formulations in function of the viscosities (η_∞_) of the different oil mixtures: (a) castor oil / MCT and (b) VEA / MCT. Different nano-emulsion size range was compared to each other and compared with pure oil mixture. FI intensity was normalized to the oil amounts in the formulations; n = 3.

In the same way as the results of Fig. 2 (b), where the FI values of two oil blends were compared, those of the nano-emulsion formulations appear very similar, even though the viscosity ranges are significantly different. The main remarkable observation lies in the fact that nano-emulsions present a higher brightness when oil is formulated compared to pure oil. In addition, FI increases when SOR increases, one can see the FI increases when the viscosity increases, but also that both *(i)* the nano-emulsification and *(ii)* the size of the droplets, have a significant impact on the fluorescence intensity. These results gather those given in Fig. 2 (b), for which the fluorescence intensity depends on the nature of the oils, even for a similar viscosity. The correlation between FI and size could be due to the modification of the slight oil composition when the presence of surfactants is gradually increased in the nano-emulsion oil core.

## 4. Conclusion

In this study, we synthesized a lipophilic fluorescent viscosity probe based on BODIPY, namely rBDP-Toco, through the coupling of a well-established molecular rotor with tocopherol. rBDP-Toco was then used to sense the viscosity of the oil core in nano-formulations such as nano-emulsions.

To date, micro-viscosity in colloidal systems is mostly measured using nano-particles tracking, and following –for instance– their magnetic properties [16], or tracking their mobility [17]. These approaches are powerful, but are limited by the size of these nanoparticles. On the other hand, molecular rotors are known to be efficient as sensor of microenvironments, and particularly lipophilic BODIPY rotors are recognized as viscosity sensor with a weak sensitivity to polarity [1,42,46]. This study is the first which proposes a deep and innovative use of lipophilic molecular rotors to characterize the viscosity in colloidal systems, reaching nanoscale compartment in the core of nano-emulsions.

We showed this experimental approach is not only efficient to sense the viscosity of lipid nano-dispersion like nano-emulsion, but also it allows to disclose specific modifications of the droplet composition, like evidence of integration of surfactants in the composition of the oil core, never identified before. The comprehensive optical calibration in oil mixtures with different viscosities was assessed and gave the correlation between the fluorescence intensity, the fluorescence lifetime, and the dynamic viscosity. It allows disclosing a universal behavior between viscosity and fluorescent lifetime, at constant temperature regardless of the composition of the oil phase. This calibration was then applied to different types of nano-emulsions encapsulating rBDP-Toco. Thus, the possibility to sense the real viscosity of the nano-droplets’ oil core, independently of their size and composition, is one of the main innovations proposed herein. In comparison with literature, mainly devoted to the use of molecular rotors for very specific targets and generally hydrophilic domains [13–15], here we propose a real calibration in lipid medium, and an innovative application to sense the properties of lipid nano-domains. In addition, this approach could also be applied to generate nano-emulsions specific micro-viscosity, by measuring and adjusting it at macroscopic scale (with oil ratio), and then control the one of nano-emulsions after formulation.

This approach allows disclosing slight variation of the oil viscosities, likely due to the presence of nonionic surfactants that remain solubilized in the oil core. A deeper characterization of the oil / surfactant mixture revealed the potential of this approach to understand the real composition of the nano-droplets. An extension of these findings could involve deploying a series of molecular motors to map the observed variations in viscosity within a given lipophilic medium, as monitored through Fluorescence Lifetime Imaging Microscopy (FLIM). Herein we establish not only the proof of the concept of the viscosity sensing inside droplets at the nanometric scale, but also shows that from slight modifications of the viscosities observed, we can identify important differences in the interactions between the compounds used in the formulation.

## Acknowledgments

M. E. acknowledges the Pierre Fabre Foundation for financial support.

## SUPPLEMENTARY INFORMATION

### 1. Synthesis of BODIPY rotor acid

BODIPY rotor acid was synthesized according to previous report [1]

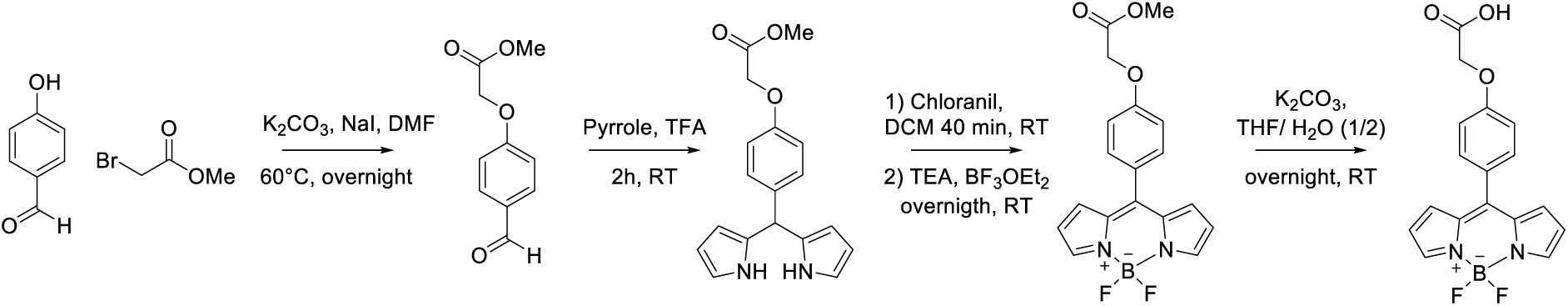

[1] Ashokkumar, P.; Ashoka, A. H.; Collot, M.; Das, A. A fluorogenic BODIPY molecular rotor as an apoptosis marker, *Chem. Commun*., **2019**, 55, 6902-6905.

**^1^H NMR (400 MHz, CD_3_OD-CDCl_3 1_-_1_)** δ 7.82 (s, 2H, H pyrrole), 7.43 (d, *J* = 8.8 Hz, 2H, H phenyl), 7.01 (d, *J* = 8.7 Hz, 2H, H phenyl), 6.87 (d, *J* = 4.2 Hz, 2H, H pyrrole), 6.48 (dd, *J* = 4.3, 2.0 Hz, 2H, H pyrrole), 4.66 (s, 2H, CH_2_).

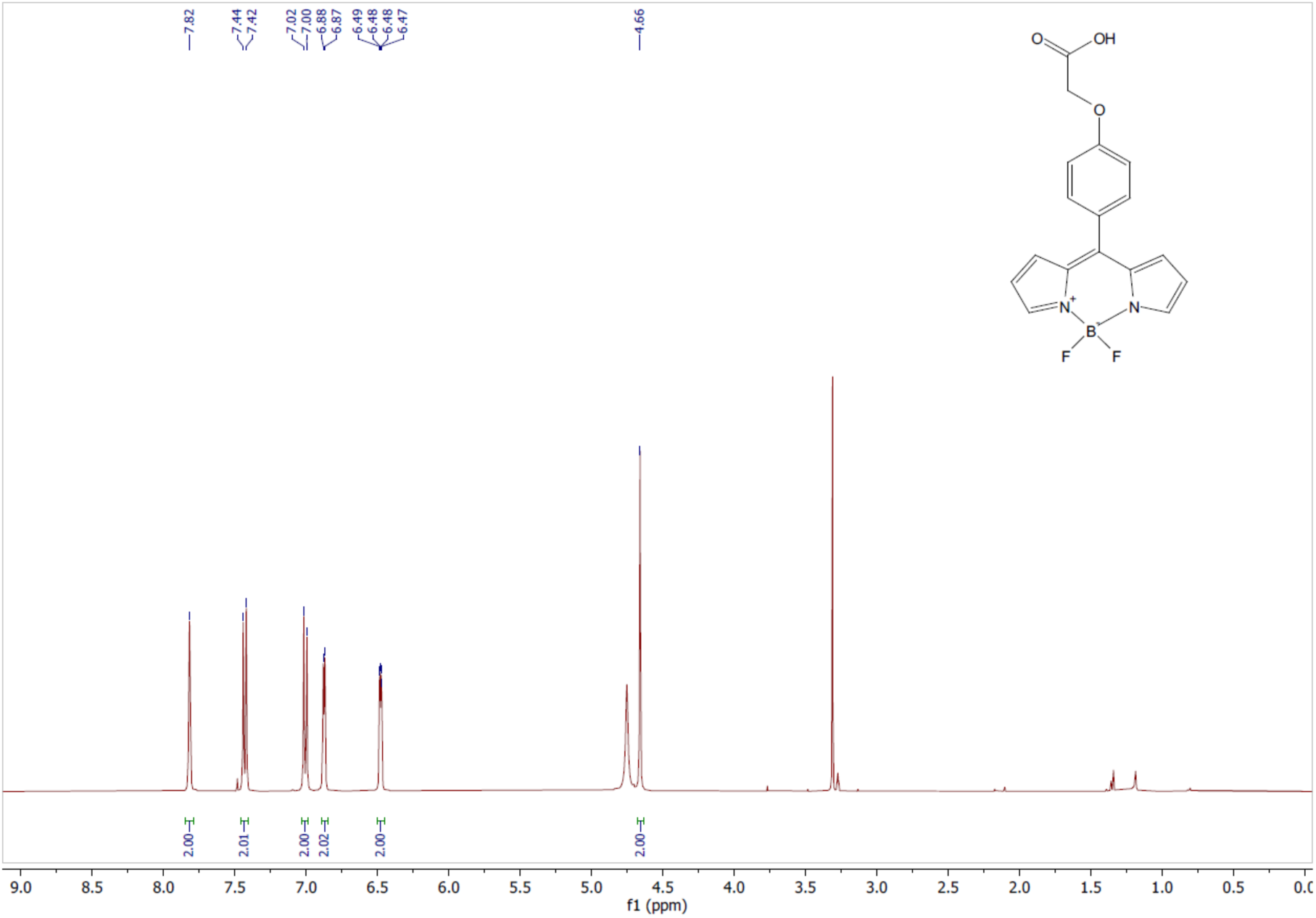

### 2. Synthesis of BODIPY rotor tocopherol

To the solution of bodipy-acid (173 mg, 0.5 mmol, 1 equiv.) dissolved in dry DCM (5 mL) was added (R)-2,5,7,8-tetramethyl-2-((4R,8R)-4,8,12-trimethyltridecyl)chroman-6-ol (322 mg, 0.75 mmol, 1.5 1 equiv.), DCC (124 mg, 0.6 mmol, 1.2 equiv.) and DMAP (12 mg, 0.1 mmol, 0.2 equiv.). The reaction mixture was stirred at room temperature overnight. The mixture was diluted with DCM and washed with water and brine, dried over MgSO4, filtered, and the solvent was removed in vacuo. The residue was purified by flash column chromatography (Heptane/40-50% DCM) to give Bodipy-rotor-tocopherol as a red oil (100 mg, 0.1 mmol, 26 %).

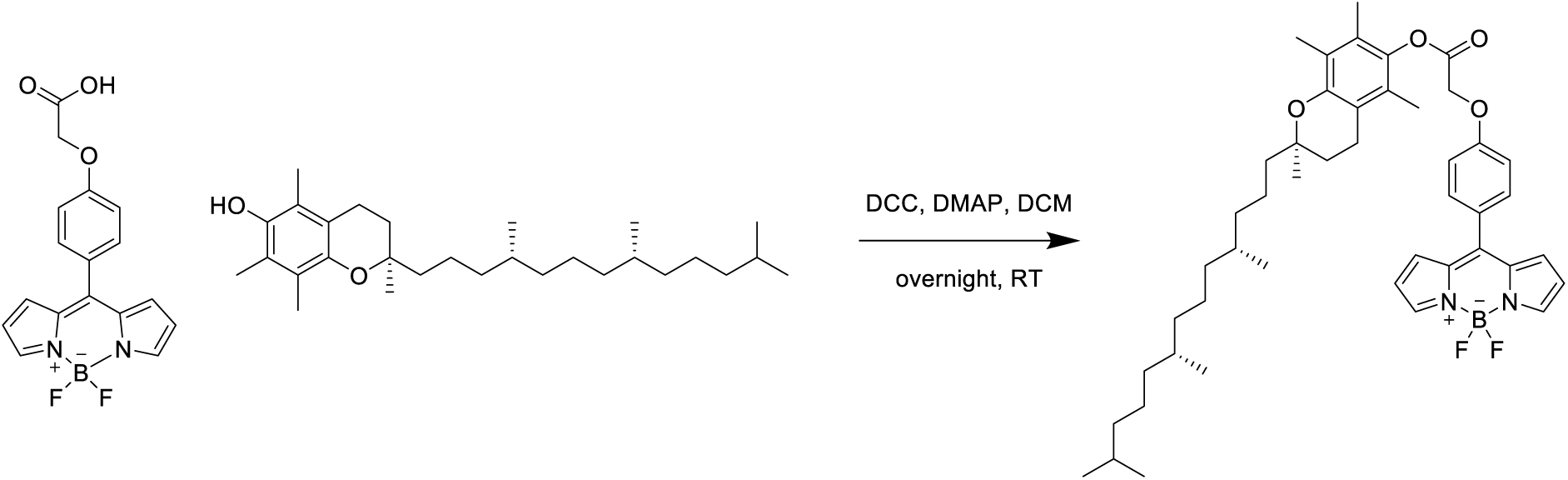

**HRMS (ESI+):** m/z calculated for C46H61BF2N2O4 M: 753,4729, found M+H: 755.4749, M+NA: 777.4569, M+K: 793.4301

**1H NMR (400 MHz, CDCl3)** δ 7.94 (s, 2H, H pyrrole), 7.58 (d, J = 8.7 Hz, 2H, H Phenyl), 7.16 (d, J = 8.8 Hz, 2H, H Phenyl), 6.96 (d, J = 4.2 Hz, 2H, H pyrrole), 6.56 (dd, J = 4.2, 1.9 Hz, 2H, H pyrrole), 5.04 (s, 2H, CH2), 2.60 (t, J = 6.8 Hz, 2H, CH2 Oxane), 2.10 (s, 3H, CH3 Benzyl), 2.02 (d, J = 18.0 Hz, 6H, CH3 Benzyl), 1.85 – 1.74 (m, 2H, CH2 Oxane), 1.60 – 1.47 (m, 4H), 1.45 – 1.35 (m, 4H, CH and CH2 aliphatic), 1.24 (m, 10H, CH3-Oxane and CH2 aliphatic), 1.16 – 1.05 (m, 6H, CH2 aliphatic), 0.89 – 0.83 (m, 12H, CH3 aliphatic).

**13C NMR (126 MHz, CDCl3)** δ 167.08 (C=O), 160.16, 149.83 (C-O), 146.92 (C-N), 143.74 (C=C-O), 139.89, 134.85, 132.39, 131.37, 127.49, 126.43, 124.74, 123.39, 118.48, 118.45, 118.42, 117.67, 114.86 (C=C), 75.25 (Cq), 65.15 (CH2), 49.11, 39.4, 37.56, 37.47, 37.42, 37.38, 37.31, 32.81, 32.80, 32.73, 32.71, 31.03, 28.00, 24.98, 24.47, 22.75, 22.66, 21.05, 20.63, (CH et CH2 aliphatic) 19.77, 19.67, 19.64 (CH3 aliphatic), 13.06, 12.21, 11.88 (CH3-Benzyl).

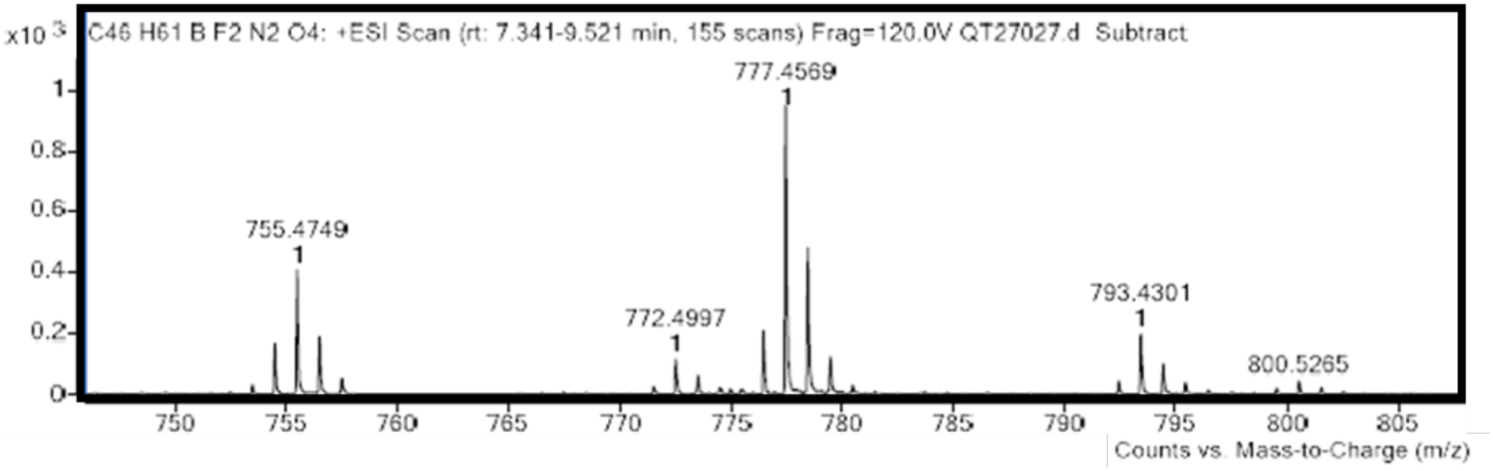

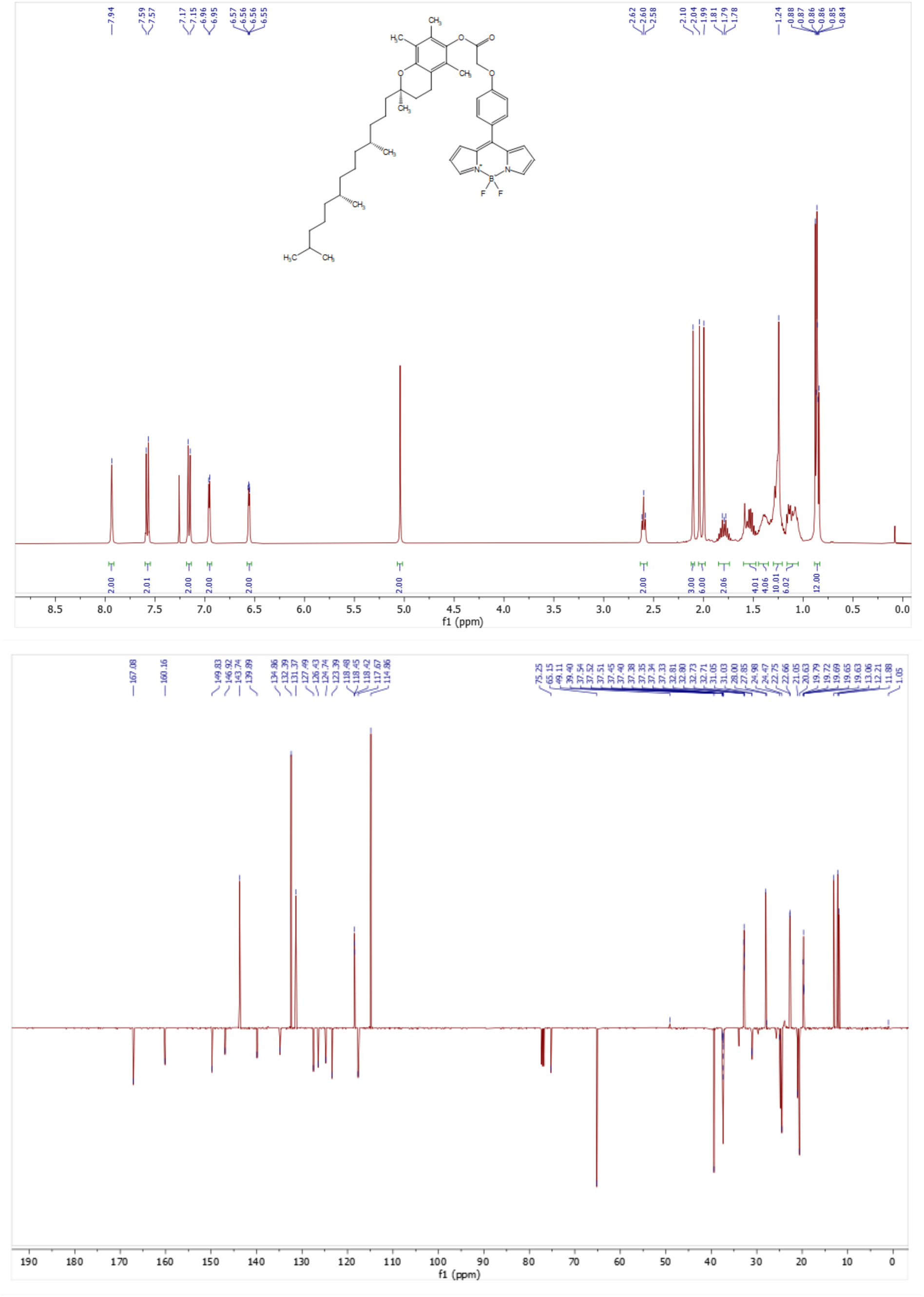

### 3. Spectral characterization of BODIPY rotor tocopherol

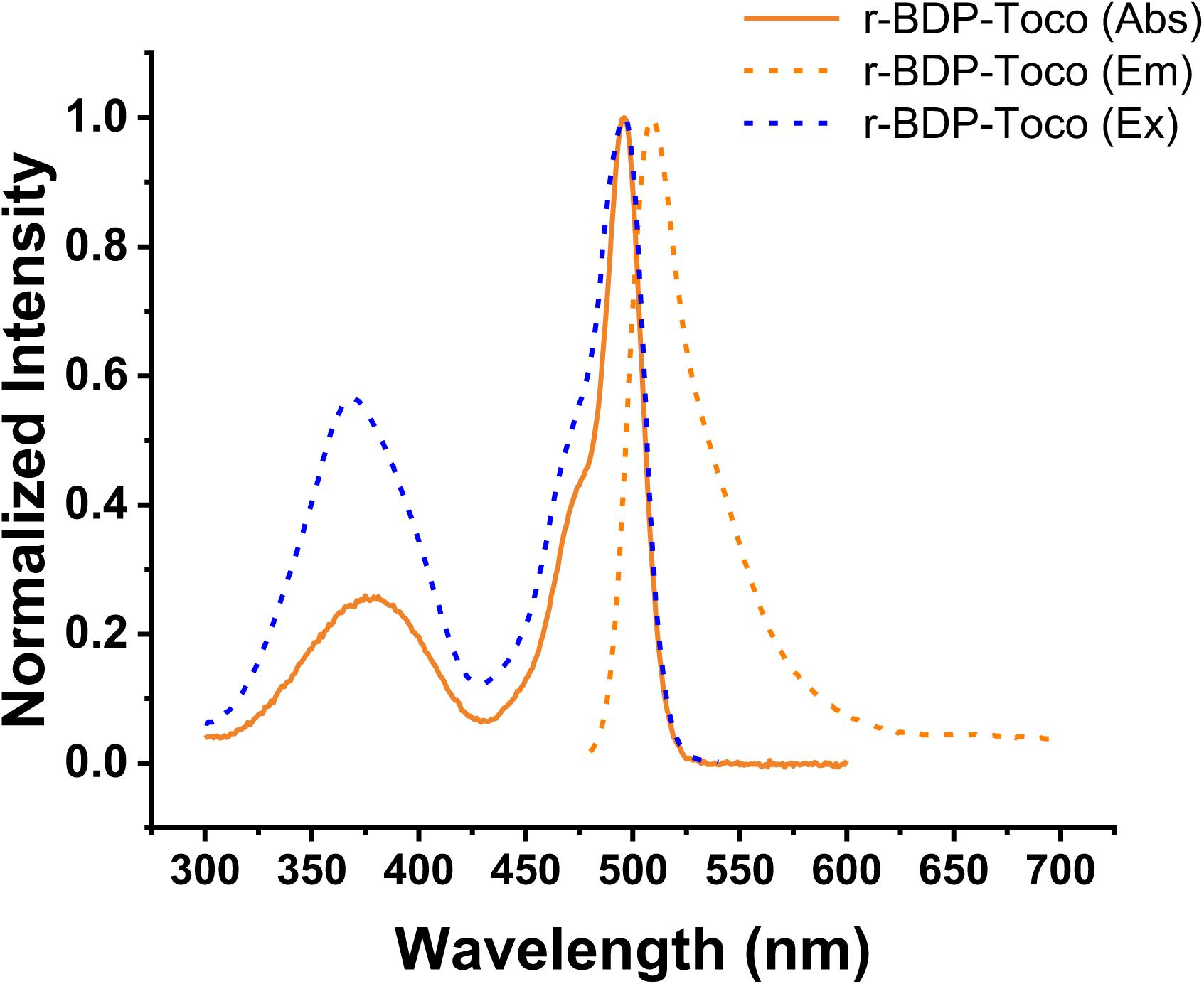

**Figure S1:**
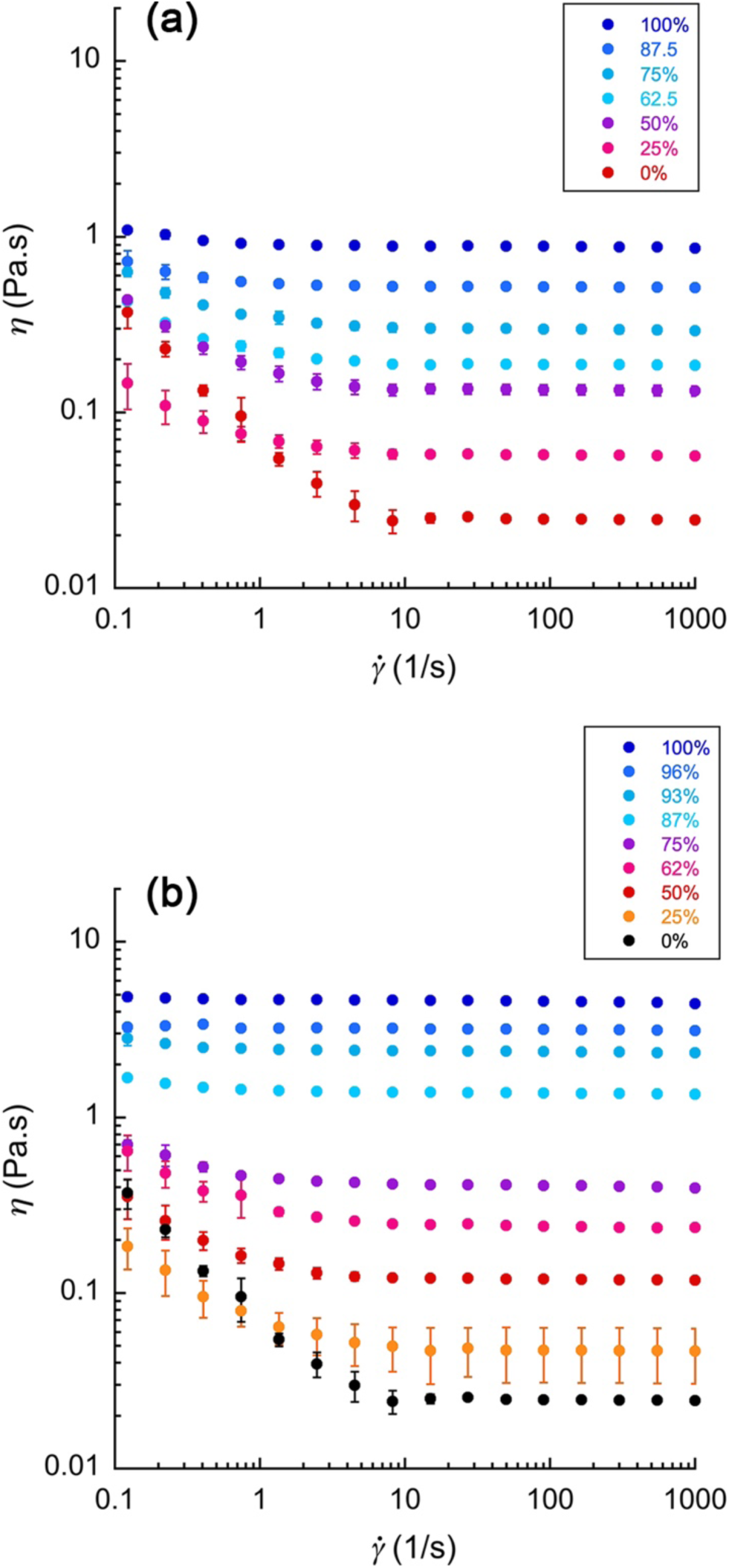
Dynamic viscosity (η) as a function of shear rate (γ̇) obtained using an oscillatory rheometer, for different percentage of (a) castor oil, and (b) VEA, mixed with MCT.

**Figure S2:**
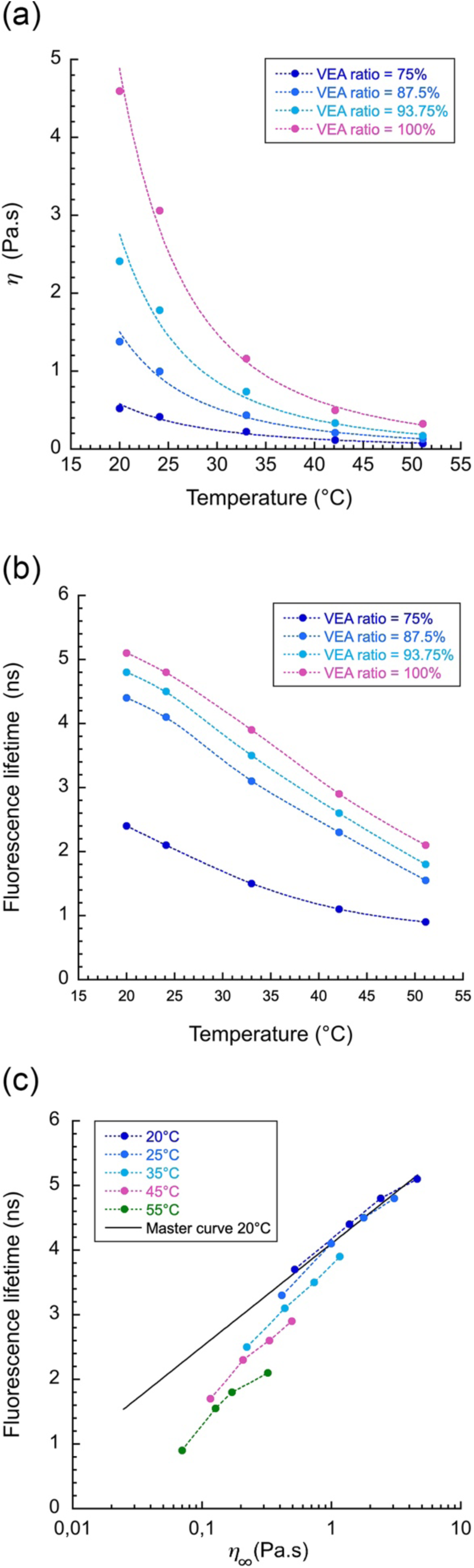
Study of the effect of temperature on rBDP-Toco molecular rotors, for VEA / MCT mixtures. (a) impact on viscosity, (b) impact on FLT, and (c) combination representing the relationship between viscosity and FLT and compared to the master curve at 20°C from Fig. 3.

**Figure S2:**
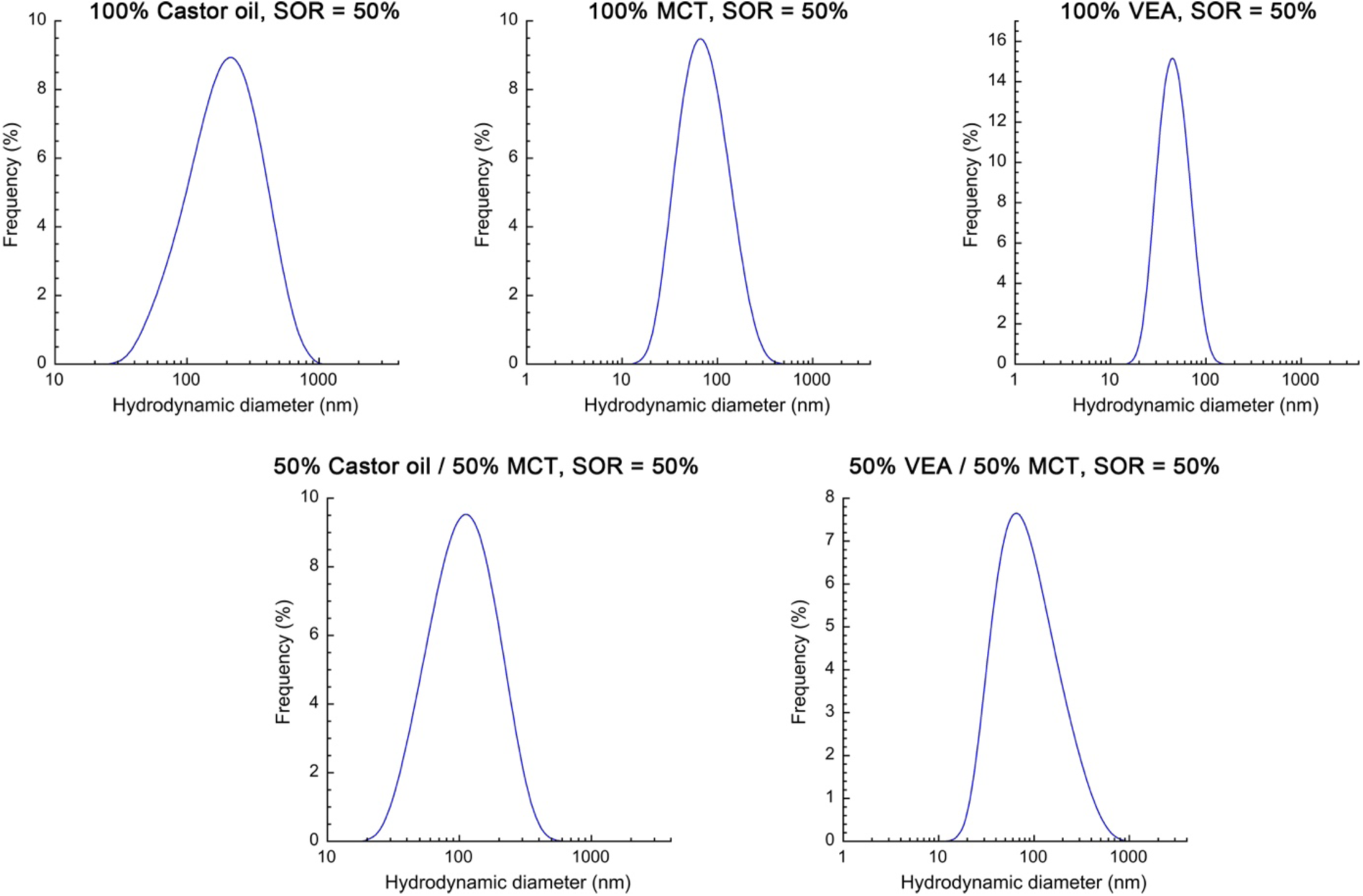
Size distribution of nano-emulsions obtained by DLS, for a panel of representative formulations.

## References

[1] A. Vyšniauskas, I. López-Duarte, N. Duchemin, T.-T. Vu, Y. Wu, E.M. Budynina, Y.A. Volkova, E. Peña Cabrera, D.E. Ramírez-Ornelas, M.K. Kuimova, Exploring viscosity, polarity and temperature sensitivity of BODIPY-based molecular rotors, Phys. Chem. Chem. Phys. 19 (2017) 25252–25259. 10.1039/C7CP03571C.

[2] M.K. Kuimova, Mapping viscosity in cells using molecular rotors, Phys. Chem. Chem. Phys. 14 (2012) 12671. 10.1039/c2cp41674c.

[3] X. Qin, X. Yang, L. Du, M. Li, Polarity-based fluorescence probes: properties and applications, RSC Med. Chem. 12 (2021) 1826–1838. 10.1039/D1MD00170A.

[4] E. Xochitiotzi-Flores, A. Jiménez-Sánchez, H. García-Ortega, N. Sánchez-Puig, M. Romero-Ávila, R. Santillan, N. Farfán, Optical properties of two fluorene derived BODIPY molecular rotors as fluorescent ratiometric viscosity probes, New J. Chem. 40 (2016) 4500–4512. 10.1039/C5NJ03339J.

[5] Z. Yang, J. Cao, Y. He, J.H. Yang, T. Kim, X. Peng, J.S. Kim, Macro-/micro-environment-sensitive chemosensing and biological imaging, Chem. Soc. Rev. 43 (2014) 4563–4601. 10.1039/C4CS00051J.

[6] J.M. Nölle, C. Jüngst, A. Zumbusch, D. Wöll, Monitoring of viscosity changes during free radical polymerization using fluorescence lifetime measurements, Polym. Chem. 5 (2014) 2700–2703. 10.1039/C3PY01684F.

[7] A. Athanasiadis, C. Fitzgerald, N.M. Davidson, C. Giorio, S.W. Botchway, A.D. Ward, M. Kalberer, F.D. Pope, M.K. Kuimova, Dynamic viscosity mapping of the oxidation of squalene aerosol particles, Phys. Chem. Chem. Phys. 18 (2016) 30385–30393. 10.1039/C6CP05674A.

[8] M. Paez-Perez, M.K. Kuimova, Molecular Rotors: Fluorescent Sensors for Microviscosity and Conformation of Biomolecules, Angew Chem Int Ed 63 (2024) e202311233. 10.1002/anie.202311233.

[9] A. Vyšniauskas, M. Qurashi, M.K. Kuimova, A Molecular Rotor that Measures Dynamic Changes of Lipid Bilayer Viscosity Caused by Oxidative Stress, Chemistry A European J 22 (2016) 13210–13217. 10.1002/chem.201601925.

[10] M.R. Dent, I. López-Duarte, C.J. Dickson, N.D. Geoghegan, J.M. Cooper, I.R. Gould, R. Krams, J.A. Bull, N.J. Brooks, M.K. Kuimova, Imaging phase separation in model lipid membranes through the use of BODIPY based molecular rotors, Phys. Chem. Chem. Phys. 17 (2015) 18393– 18402. 10.1039/C5CP01937K.

[11] E. Gatzogiannis, Z. Chen, L. Wei, R. Wombacher, Y.-T. Kao, G. Yefremov, V.W. Cornish, W. Min, Mapping protein-specific micro-environments in live cells by fluorescence lifetime imaging of a hybrid genetic-chemical molecular rotor tag, Chem. Commun. 48 (2012) 8694. 10.1039/c2cc33133k.

[12] X. Peng, Z. Yang, J. Wang, J. Fan, Y. He, F. Song, B. Wang, S. Sun, J. Qu, J. Qi, M. Yan, Fluorescence Ratiometry and Fluorescence Lifetime Imaging: Using a Single Molecular Sensor for Dual Mode Imaging of Cellular Viscosity, J. Am. Chem. Soc. 133 (2011) 6626–6635. 10.1021/ja1104014.

[13] A. Vyšniauskas, M.K. Kuimova, A twisted tale: measuring viscosity and temperature of microenvironments using molecular rotors, International Reviews in Physical Chemistry 37 (2018) 259–285. 10.1080/0144235X.2018.1510461.

[14] S. Toliautas, J. Dodonova, A. Žvirblis, I. Čiplys, A. Polita, A. Devižis, S. Tumkevičius, J. Šulskus, A. Vyšniauskas, Enhancing the Viscosity-Sensitive Range of a BODIPY Molecular Rotor by Two Orders of Magnitude, Chemistry A European J 25 (2019) 10342–10349. 10.1002/chem.201901315.

[15] L. Michels, V. Gorelova, Y. Harnvanichvech, J.W. Borst, B. Albada, D. Weijers, J. Sprakel, Complete microviscosity maps of living plant cells and tissues with a toolbox of targeting mechanoprobes, Proc. Natl. Acad. Sci. U.S.A. 117 (2020) 18110–18118. 10.1073/pnas.1921374117.

[16] R.R. Merchant, L. Maldonado-Camargo, C. Rinaldi, In situ measurements of dispersed and continuous phase viscosities of emulsions using nanoparticles, Journal of Colloid and Interface Science 486 (2017) 241–248. 10.1016/j.jcis.2016.09.063.

[17] T. Moschakis, A. Lazaridou, C.G. Biliaderis, Using particle tracking to probe the local dynamics of barley β-glucan solutions upon gelation, Journal of Colloid and Interface Science 375 (2012) 50–59. 10.1016/j.jcis.2012.02.048.

[18] B. Medronho, A. Filipe, C. Costa, A. Romano, B. Lindman, H. Edlund, M. Norgren, Microrheology of novel cellulose stabilized oil-in-water emulsions, Journal of Colloid and Interface Science 531 (2018) 225–232. 10.1016/j.jcis.2018.07.043.

[19] X. Li, J. van der Gucht, P. Erni, R. de Vries, Active microrheology of protein condensates using colloidal probe-AFM, Journal of Colloid and Interface Science 632 (2023) 357–366. 10.1016/j.jcis.2022.11.071.

[20] S.O. Raja, G. Sivaraman, S. Biswas, G. Singh, F. Kalim, P. Kandaswamy, A. Gulyani, A Tunable Palette of Molecular Rotors Allows Multicolor, Ratiometric Fluorescence Imaging and Direct Mapping of Mitochondrial Heterogeneity, ACS Appl. Bio Mater. 4 (2021) 4361–4372. 10.1021/acsabm.1c00135.

[21] X. Wu, C. Barner-Kowollik, Fluorescence-readout as a powerful macromolecular characterisation tool, Chem. Sci. 14 (2023) 12815–12849. 10.1039/D3SC04052F.

[22] M.Y. Berezin, S. Achilefu, Fluorescence Lifetime Measurements and Biological Imaging, Chem. Rev. 110 (2010) 2641–2684. 10.1021/cr900343z.

[23] A. Polita, S. Toliautas, R. Žvirblis, A. Vyšniauskas, The effect of solvent polarity and macromolecular crowding on the viscosity sensitivity of a molecular rotor BODIPY-C 10, Phys. Chem. Chem. Phys. 22 (2020) 8296–8303. 10.1039/C9CP06865A.

[24] T.T. Vu, R. Méallet-Renault, G. Clavier, B.A. Trofimov, M.K. Kuimova, Tuning BODIPY molecular rotors into the red: sensitivity to viscosity vs. temperature, J. Mater. Chem. C 4 (2016) 2828–2833. 10.1039/C5TC02954F.

[25] T.F. Vandamme, The universality of low-energy nano-emulsification, International Journal of Pharmaceutics 377 (2009) 142–147. 10.1016/j.ijpharm.2009.05.014.

[26] M. Yao, H. Xiao, D.J. McClements, Delivery of Lipophilic Bioactives: Assembly, Disassembly, and Reassembly of Lipid Nanoparticles, Annu. Rev. Food Sci. Technol. 5 (2014) 53–81. 10.1146/annurev-food-072913-100350.

[27] V.K. Rai, N. Mishra, K.S. Yadav, N.P. Yadav, Nanoemulsion as pharmaceutical carrier for dermal and transdermal drug delivery: Formulation development, stability issues, basic considerations and applications, Journal of Controlled Release 270 (2018) 203–225. 10.1016/j.jconrel.2017.11.049.

[28] M. Kah, T. Hofmann, Nanopesticide research: Current trends and future priorities, Environment International 63 (2014) 224–235. 10.1016/j.envint.2013.11.015.

[29] Ş. Yalçınöz, E. Erçelebi, Potential applications of nano-emulsions in the food systems: an update, Mater. Res. Express 5 (2018) 062001. 10.1088/2053-1591/aac7ee.

[30] M.N. Yukuyama, D.D.M. Ghisleni, T.J.A. Pinto, N.A. Bou-Chacra, Nanoemulsion: process selection and application in cosmetics – a review, Intern J of Cosmetic Sci 38 (2016) 13–24. 10.1111/ics.12260.

[31] A.S. Klymchenko, F. Liu, M. Collot, N. Anton, Dye-Loaded Nanoemulsions: Biomimetic Fluorescent Nanocarriers for Bioimaging and Nanomedicine, Adv. Healthcare Mater. 10 (2021) 2001289. 10.1002/adhm.202001289.

[32] X. Li, N. Anton, G. Zuber, T. Vandamme, Contrast agents for preclinical targeted X-ray imaging, Advanced Drug Delivery Reviews 76 (2014) 116–133. 10.1016/j.addr.2014.07.013.

[33] N. Anton, F. Hallouard, M.F. Attia, T.F. Vandamme, Nano-emulsions for Drug Delivery and Biomedical Imaging, in: A. Prokop, V. Weissig (Eds.), Intracellular Delivery III, Springer International Publishing, Cham, 2016: pp. 273–300. 10.1007/978-3-319-43525-1_11.

[34] A.S. Klymchenko, E. Roger, N. Anton, H. Anton, I. Shulov, J. Vermot, Y. Mely, T.F. Vandamme, Highly lipophilic fluorescent dyes in nano-emulsions: towards bright non-leaking nano-droplets, RSC Adv. 2 (2012) 11876. 10.1039/c2ra21544f.

[35] V.N. Kilin, H. Anton, N. Anton, E. Steed, J. Vermot, T.F. Vandamme, Y. Mely, A.S. Klymchenko, Counterion-enhanced cyanine dye loading into lipid nano-droplets for single-particle tracking in zebrafish, Biomaterials 35 (2014) 4950–4957. 10.1016/j.biomaterials.2014.02.053.

[36] X. Wang, N. Anton, P. Ashokkumar, H. Anton, T.K. Fam, T. Vandamme, A.S. Klymchenko, M. Collot, Optimizing the Fluorescence Properties of Nanoemulsions for Single Particle Tracking in Live Cells, ACS Appl. Mater. Interfaces 11 (2019) 13079–13090. 10.1021/acsami.8b22297.

[37] X. Ma, R. Sun, J. Cheng, J. Liu, F. Gou, H. Xiang, X. Zhou, Fluorescence Aggregation-Caused Quenching versus Aggregation-Induced Emission: A Visual Teaching Technology for Undergraduate Chemistry Students, J. Chem. Educ. 93 (2016) 345–350. 10.1021/acs.jchemed.5b00483.

[38] A. Mushtaq, S. Mohd Wani, A.R. Malik, A. Gull, S. Ramniwas, G. Ahmad Nayik, S. Ercisli, R. Alina Marc, R. Ullah, A. Bari, Recent insights into Nanoemulsions: Their preparation, properties and applications, Food Chemistry: X 18 (2023) 100684. 10.1016/j.fochx.2023.100684.

[39] D. Myers, Liquid–Fluid Interfaces, in: Surfaces, Interfaces, and Colloids, John Wiley & Sons, Ltd, n.d.: pp. 140–178. 10.1002/0471234990.ch8.

[40] W.D. Bancroft, The Theory of Emulsification, V, J. Phys. Chem. 17 (1913) 501–519. 10.1021/j150141a002.

[41] W. Miao, C. Yu, E. Hao, L. Jiao, Functionalized BODIPYs as Fluorescent Molecular Rotors for Viscosity Detection, Front. Chem. 7 (2019) 825. 10.3389/fchem.2019.00825.

[42] X. Wang, S. Bou, A.S. Klymchenko, N. Anton, M. Collot, Ultrabright Green-Emitting Nanoemulsions Based on Natural Lipids-BODIPY Conjugates, Nanomaterials 11 (2021). 10.3390/nano11030826.

[43] P. Ashokkumar, A.H. Ashoka, M. Collot, A. Das, A.S. Klymchenko, A fluorogenic BODIPY molecular rotor as an apoptosis marker, Chem. Commun. 55 (2019) 6902–6905. 10.1039/C9CC03242H.

[44] K.T. Fam, L. Saladin, A.S. Klymchenko, M. Collot, Confronting molecular rotors and self-quenched dimers as fluorogenic BODIPY systems to probe biotin receptors in cancer cells, Chem. Commun. 57 (2021) 4807–4810. 10.1039/D1CC00108F.

[45] Y. Wu, M. Štefl, A. Olzyńska, M. Hof, G. Yahioglu, P. Yip, D.R. Casey, O. Ces, J. Humpolíčková, M.K. Kuimova, Molecular rheometry: direct determination of viscosity in Lo and Ld lipid phases via fluorescence lifetime imaging, Phys. Chem. Chem. Phys. 15 (2013) 14986–14993. 10.1039/C3CP51953H.

[46] J.A. Levitt, P.-H. Chung, M.K. Kuimova, G. Yahioglu, Y. Wang, J. Qu, K. Suhling, Fluorescence Anisotropy of Molecular Rotors, ChemPhysChem 12 (2011) 662–672. 10.1002/cphc.201000782.

[47] A. Vyšniauskas, M. Qurashi, N. Gallop, M. Balaz, H.L. Anderson, M.K. Kuimova, Unravelling the effect of temperature on viscosity-sensitive fluorescent molecular rotors, Chem. Sci. 6 (2015) 5773–5778. 10.1039/C5SC02248G.

[48] A.U. Rehman, Spontaneous nano-emulsification with tailor-made amphiphilic polymers and related monomers, Eur. J. Pharm. Res. 1 (2019) 27–36. 10.34154/2019-EJPR.01(01).pp-27-36/euraass.

[49] S. Ding, B. Mustafa, N. Anton, C.A. Serra, D. Chan-Seng, T.F. Vandamme, Production of lipophilic nanogels by spontaneous emulsification, International Journal of Pharmaceutics 585 (2020) 119481. 10.1016/j.ijpharm.2020.119481.

[50] J. Stetefeld, S.A. McKenna, T.R. Patel, Dynamic light scattering: a practical guide and applications in biomedical sciences, Biophysical Reviews 8 (2016) 409–427. 10.1007/s12551-016-0218-6.

[51] M. Zhang, Y. Yang, N.C. Acevedo, Effect of Oil Content and Composition on the Gelling Properties of Egg-SPI Proteins Stabilized Emulsion Gels, Food Biophysics 15 (2020) 473–481. 10.1007/s11483-020-09646-8.

[52] N. Anton, T.F. Vandamme, The universality of low-energy nano-emulsification, International Journal of Pharmaceutics 377 (2009) 142–147. 10.1016/j.ijpharm.2009.05.014.

[53] S. Akram, N. Anton, Z. Omran, T. Vandamme, Water-in-Oil Nano-Emulsions Prepared by Spontaneous Emulsification: New Insights on the Formulation Process, Pharmaceutics 13 (2021) 1030. 10.3390/pharmaceutics13071030.

[54] X. Wang, M. Collot, T.F. Vandamme, N. Anton, Study of the spontaneous nano-emulsification process with different octadecyl succinic anhydride derivatives, Colloids and Surfaces A: Physicochemical and Engineering Aspects 645 (2022) 128858. 10.1016/j.colsurfa.2022.128858.

[55] M.F. Attia, N. Anton, R. Akasov, M. Chiper, E. Markvicheva, T.F. Vandamme, Biodistribution and Toxicity of X-Ray Iodinated Contrast Agent in Nano-emulsions in Function of Their Size, Pharmaceutical Research 33 (2016) 603–614. 10.1007/s11095-015-1813-0.

[56] M.F. Attia, N. Anton, M. Chiper, R. Akasov, H. Anton, N. Messaddeq, S. Fournel, A.S. Klymchenko, Y. Mély, T.F. Vandamme, Biodistribution of X-Ray Iodinated Contrast Agent in Nano-Emulsions Is Controlled by the Chemical Nature of the Oily Core, ACS Nano 8 (2014) 10537–10550. 10.1021/nn503973z.

[57] A.H. Saberi, Y. Fang, D.J. McClements, Fabrication of vitamin E-enriched nanoemulsions: Factors affecting particle size using spontaneous emulsification, Journal of Colloid and Interface Science 391 (2013) 95–102. 10.1016/j.jcis.2012.08.069.

